# Novel Complete Methanogenic Pathways in Longitudinal Genomic Study of Monogastric Age-Associated Archaea

**DOI:** 10.1101/2022.12.03.518977

**Authors:** Brandi Feehan, Qinghong Ran, Victoria Dorman, Kourtney Rumback, Sophia Pogranichniy, Kaitlyn Ward, Robert Goodband, Megan C Niederwerder, Sonny T M Lee

## Abstract

**Background:** Archaea perform critical roles in the microbiome system, including utilizing hydrogen to allow for enhanced microbiome member growth and influencing overall host health. With the majority of microbiome research focussing on bacteria, the functions of archaea are largely still under investigation. Understanding methanogenic functions during the host lifetime will add to the limited knowledge on archaeal influence on gut and host health. In our study, we determined lifelong archaea detection and methanogenic functions while assessing global and host distribution of our novel archaeal metagenome assembled genomes (MAGs). We followed 7 monogastric swine throughout their life, from birth to adult (1-156 days of age), and collected feces at 22 time points. The samples underwent gDNA extraction, Illumina sequencing, bioinformatic quality and assembly processes, and MAG taxonomic assignment and functional annotation.

**Results:** We generated 1,130 non-redundant MAGs with 8 classified as methanogenic archaea. The taxonomic classifications were as follows: orders *Methanomassiliicoccales* (5) and *Methanobacteriales* (3); genera *UBA71* (3), *Methanomethylophilus* (1), *MX-02* (1), and *Methanobrevibacter* (3). We recovered the first US swine *Methanobrevibacter UBA71 sp006954425* and *Methanobrevibacter gottschalkii* MAGs. The *Methanobacteriales* MAGs were identified primarily during the young, preweaned host whereas *Methanomassiliicoccales* primarily in the adult host. Moreover, we identified our methanogens in metagenomic sequences from Chinese swine, US adult humans, Mexican adult humans, Swedish adult humans, and paleontological humans, indicating that methanogens span different hosts, geography and time. We determined complete metabolic pathways for all three methanogenic pathways: hydrogenotrophic, methylotrophic, and acetoclastic. This study provided the first evidence of acetoclastic methanogenesis in monogastric archaea which indicated a previously unknown capability for acetate utilization in methanogenesis for monogastric methanogens. Overall, we hypothesized that the age-associated detection patterns were due to differential substrate availability via the host diet and microbial metabolism, and that these methanogenic functions are likely crucial to methanogens across hosts. This study provided a comprehensive, genome-centric investigation of monogastric-associated methanogens which will further our understanding of microbiome development and functions.

## Introduction

The gastrointestinal system contains countless microorganisms spanning multiple kingdoms performing equally diverse functions. Archaea, bacteria, viruses, and fungi work in concert and competition to acquire nutrients and space^1^. The focus of previous gut microbiome research has predominantly been on the identification and function of bacteria^2, 3^. However, archaea have been demonstrated to be equally important members of the gastrointestinal microbiome^4^. Methanogenic archaea, or archaea which carry out methanogenesis, perform crucial roles in the gut^4, 5^. Yet, current research has not indicated how methanogenic gut function changes throughout the lifetime of monogastric hosts^6, 7^. With limited research on archaea, and even more minimal analysis on methanogenic functions, we are lacking an in-depth understanding of gastrointestinal associated methanogens, especially our comprehension of methanogen influence on gut and host health throughout host stages of life. By investigating monogastric associated methanogens with a longitudinal approach, we are adding essential knowledge to the limited understanding of monogastric methanogens.

While some beneficial and detrimental associations of archaea to host health have been reported, overall the role of archaea in health and disease is still under investigation^5^. To date, archaea have been associated with a few illnesses including: brain abscesses^5, 8^, sinus abscesses^5, 9^, and several gastrointestinal disorders, such as constipation^5, 10, 11^ and obesity^5, 12^. Conversely, archaea have also been associated with beneficial attributes. For example, archaea metabolize trimethylamine (TMA), which is thought to decrease cardiovascular disease^4, 5^, has prompted further evaluation of archaea members as a probiotic for cardiovascular health^5, 13^. Moreover, archaea allow continued microbial metabolism, growth and action by lowering hydrogen gut levels^5^. Archaea’s role of hydrogen utilization is especially important in the gut where microorganisms work in concert within the shared gut-microbiome system. However with limited prior research, there is a critical need to understand the role of gastrointestinal archaea in health and sickness via hydrogen metabolism.

Overall, archaea are classified into four superphyla: Euryarchaeota, Asgard, TACK (Thaumarchaeota, Aigarchaeota, Crenarchaeota and Korarchaeota), and DPANN (Diapherotrites, Parvarchaeota, Aenigmarchaeota, Nanoarchaeota, and Nanohaloarchaeota)^14^. To date, Asgard archaea have not been indicated as methanogens^14^, and TACK and DPANN have only been identified in non-host associated environmental sites^14–16^. Therefore, currently known host-associated gut methanogens fall within the seven orders of Euryarchaeota: Methanobacteriales, Methanococcales, Methanomicrobiales, Methanosarcinales, Methanocellales, Methanopyrales, Methanomassiliicoccales^17–20^. These Euryarchaeota orders are obligate anaerobes which perform methanogenesis to conserve energy for ATP production, where methane is a byproduct^17, 21^. Actions immediately following methanogenesis generate an ion gradient which is coupled with ATP production^22, 23^.

Given the necessity for ATP production, it is unsurprising that historically, studies have primarily relied on the methanogenic gene methyl-coenzyme M reductase A (*mcrA*) or 16S rRNA for identification of gut-associated methanogens^7, 24–26^. *McrA* has been identified in all methanogens to date, as the protein performs a critical role in the final methane production step of methanogenesis^26, 27^. While prior research was heavily reliant on targeted PCR methodologies, we are missing a functional understanding, from complete genetic sequencing, of gut-associated methanogens^28^. Functional methanogen studies become even more profound when evaluated in a longitudinal approach, especially when following the same hosts. In doing so, we can determine lifetime gut methanogen dynamics and host implications. Currently, studies which evaluate longitudinal methanogen dynamics typically involve ruminants, such as cows, sheep, goats, and deer^29^. At the time of publication, we could not find a longitudinal study of methanogen genomes (i.e. not marker studies such as 16S rRNA or *mcrA*) following the same monogastrics hosts throughout their lifetime, highlighting the crucial need for such metagenomic longitudinal evaluations^6, 7^. Without this knowledge, we cannot determine lifetime dynamics of archaea, and how their methanogenic function may be related to age-associated factors, such as diet and host development.

Our study is the first description of longitudinal monogastric methanogens with genomic analysis of methanogenic function. We evaluated methanogen abundance and functions of 7 monogastric swine hosts over their lifetime at 22 timepoints from birth through adulthood (ages 1-156 days). We described how our methanogen metagenome assembled genomes (MAGs) were identified during specific host ages. Furthermore, we determined methanogenic pathways previously unknown to monogastric-associated methanogens. This study provided evidence of multiple novel methanogen characteristics, which will aid future studies as we build the monogastric methanogen repertoire.

## Results and Discussion

Host-associated archaeal methanogens have been linked to various conditions of health and disease. Most archaea-centric intestinal microbiome studies have been conducted on a single time point in the lifetime of the host. Using molecular and cultural approaches, intestinal archaea have been identified in many hosts, including: humans, swine, horses, rats, birds, fish, and kangaroos^30^. Overall, these analyses reported that the most common methanogens in the gut are members of the *Methanobacteriales* and *Methanomassiliicoccales* orders^30^. However, little is known about the presence and distribution of archaea through the lifetime of the swine. There is also a lack of data on the functions of the archaea in the swine gut. Overall, this knowledge gap has hindered the identification of factors that influence the diversity, abundance and functions of archaea in the swine. In this study, we recovered 8 methanogenic archaea metagenome assembled genomes (MAGs) that exhibited differential colonization patterns in the host at different ages. While distribution of methanogens across multiple hosts has been previously demonstrated, we recovered the first US swine

### Methanobrevibacter UBA71 sp006954425 and Methanobrevibacter gottschalkii MAGs^30^

Moreover, we attributed methanogenic functions to our age-associated archaea, and identified the first evidence of acetoclastic methanogenesis in monogastric archaea, found in our *Methanomassiliicoccales* MAGs, indicating a previously unknown capability of monogastric methanogens to utilize acetate in energy acquisition. Alternatively, we attributed hydrogenotrophic methanogenesis, where carbon dioxide (CO_2_) is utilized, in the *Methanobacteriales*. We surmised that the age-associated detection patterns were due to differential substrate availability, which was highly influenced by diet. Altogether, we provided a comprehensive, genome-centric investigation of monogastric-associated archaea to further our understanding of microbiome development and function.

### Taxonomic classification of gut metagenome-assembled genomes

To broadly sample gut-associated microorganisms of the swine host across different age-associated growth stages, we obtained 5,840,640,191 paired-end reads from Illumina NovaSeq sequencing data of 112 swine fecal samples (Supplementary Table S3). After quality trimming, we generated 5,167,665,150 paired-end reads. The resulting 3 co-assemblies contained 9,431,702 contigs that described approximately ∼3.6 x 10^10^ nucleotides and ∼3.7 x 10^7^ genes. Using a combination of automatic and manual binning strategies resulted in 4,556 metagenome-assembled genomes (MAGs). We further removed redundancy by selecting a single representative for each set of genomes that shared an average nucleotide identity (ANI) of greater than 95%, resulting in 1,130 final non-redundant MAGs (nr-MAGs) (Supplementary Table S3). Among the nr-MAGs, we recovered an average of 203 ± 187 contigs, with an average N50 of 32,737 ± 35,205. The resolved nr-MAGs had completion values of 87.9% ± 8.6%. The genomic lineages for archaeal and bacterial nr-MAGs based on domain-specific single-copy core genes resolved to 20 phyla (2 archaea phyla and 18 bacterial phyla). We could also assign 88.4% of the bacterial and archaeal nr-MAGs to their genera.

### Resolved archaeal MAGs phylogenetically similar to diverse hosts and geographic disbursed archaea

Among the 1,130 nr-MAGs that we resolved, our genomic collection also included 8 archaea nr-MAGs (hereafter known as archaea-MAGs; Ar-1 through Ar-8; Table 1; Supplementary Table S3). We observed that our resolved archaea-MAGs harbored genes which encoded for critical methyl-coenzyme M reductase (*mcrABG*) proteins required for methanogenesis, including *mcrA* which is typically utilized for methanogen classification^31, 32^ (Supplementary Tables S3 and S4). To our best knowledge, these MAGs represent the first genomic evidence of putative methanogens differential colonization pattern of the monogastric gut. The resolved methanogen MAGs had an average genome size of 1.4 Mbp, 1,573 KEGG gene annotations, 1,535 COG gene annotations, and a GC content ranging from 31% to 56% (Table 1; Supplementary Table S3). We resolved 7 of the methanogen MAGs to the species level with one archaea-MAG resolving to the genus level (Table 1; Supplementary Table S3). Our resolved archaea-MAGs were assigned to the following orders: *Methanomassiliicoccales* (5) and *Methanobacteriales* (3). Moreover, the genera were as follows: *UBA71* (3), *Methanomethylophilus* (1), *MX-02* (1), and *Methanobrevibacter* (3).

**Table 1.**
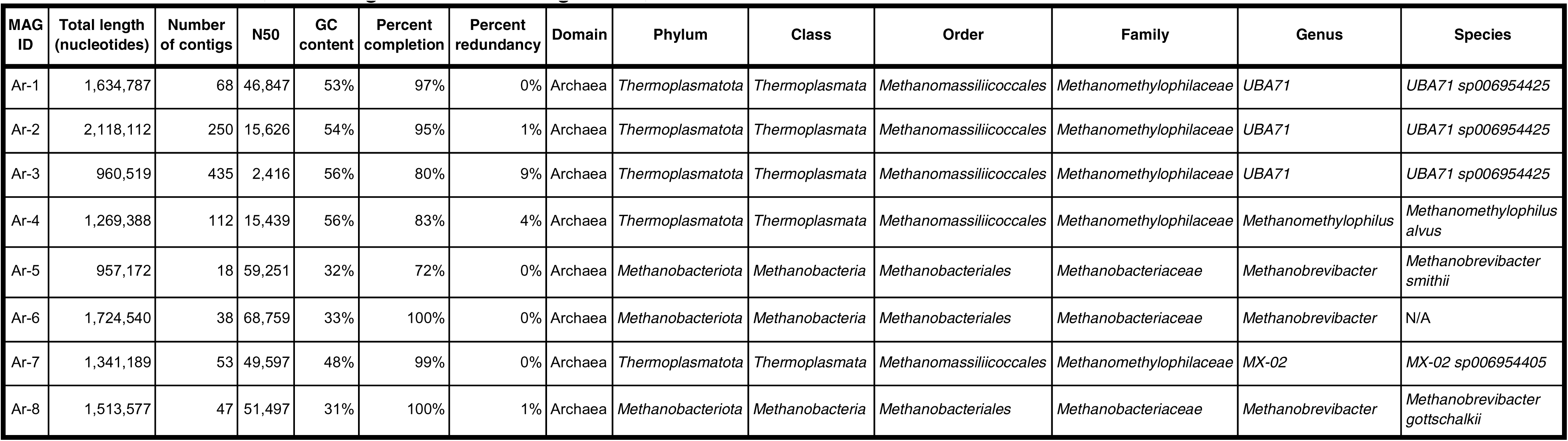
Anvi’o results, including taxonomic assignment, of 8 archaea-MAGs.

| MAG ID | Total length (nucleotides) | Number of contigs | N50 | GC content | Percent completion | Percent redundancy | Domain | Phylum | Class | Order | Family | Genus | Species |
| --- | --- | --- | --- | --- | --- | --- | --- | --- | --- | --- | --- | --- | --- |
| Ar-1 | 1,634,787 | 68 | 46,847 | 53% | 97% | 0% | Archaea | Thermoplasmatota | Thermoplasmata | Methanomassiliicoccales | Methanomethylophilaceae | UBA71 | UBA71 sp006954425 |
| Ar-2 | 2,118,112 | 250 | 15,626 | 54% | 95% | 1% | Archaea | Thermoplasmatota | Thermoplasmata | Methanomassiliicoccales | Methanomethylophilaceae | UBA71 | UBA71 sp006954425 |
| Ar-3 | 960,519 | 435 | 2,416 | 56% | 80% | 9% | Archaea | Thermoplasmatota | Thermoplasmata | Methanomassiliicoccales | Methanomethylophilaceae | UBA71 | UBA71 sp006954425 |
| Ar-4 | 1,269,388 | 112 | 15,439 | 56% | 83% | 4% | Archaea | Thermoplasmatota | Thermoplasmata | Methanomassiliicoccales | Methanomethylophilaceae | Methanomethylophilus | Methanomethylophilus alvus |
| Ar-5 | 957,172 | 18 | 59,251 | 32% | 72% | 0% | Archaea | Methanobacteriota | Methanobacteria | Methanobacteriales | Methanobacteriaceae | Methanobrevibacter | Methanobrevibacter smithii |
| Ar-6 | 1,724,540 | 38 | 68,759 | 33% | 100% | 0% | Archaea | Methanobacteriota | Methanobacteria | Methanobacteriales | Methanobacteriaceae | Methanobrevibacter | N/A |
| Ar-7 | 1,341,189 | 53 | 49,597 | 48% | 99% | 0% | Archaea | Thermoplasmatota | Thermoplasmata | Methanomassiliicoccales | Methanomethylophilaceae | MX-02 | MX-02 sp006954405 |
| Ar-8 | 1,513,577 | 47 | 51,497 | 31% | 100% | 1% | Archaea | Methanobacteriota | Methanobacteria | Methanobacteriales | Methanobacteriaceae | Methanobrevibacter | Methanobrevibacter gottschalkii |

We downloaded 95 *Methanomassiliicoccales* and 97 *Methanobacteriales* genomes to investigate the phylogenetic relationship of our resolved archaea-MAGs (Figure 2; Supplementary Figure S1). We showed that our methanogen populations had close phylogenetic relationships with archaea from geographically distinct mammalian hosts, suggesting high similarities in gene functions in archaea among diverse host species. Given similarities amongst such diverse host species with diverse digestive systems, we hypothesize these close genetic relatives of our resolved archaea-MAGs might be more ubiquitous in a wider range of hosts than are currently discussed. We noticed Ar-4 clustered, as expected, with 6 *Methanomethylophilus alvus* strains: 5 from human gut samples and 1 from swine (MAG221) (Figure 2A)^33–38^. Ar-7 clustered with 4 *MX-02 sp006954405*. United Kingdom strain 10^39^ and Chinese strain MAG014^36^ have been identified as swine-originating, whereas B5_69.fa and B45_maxbin.030.fa were from humans^40^. Interestingly, clustering on the same branch (*B5_69.fa* and *B45_maxbin.030.fa*) are archaea isolated from South African adult humans^40^. Ar-1 was in the same branch with archaea isolated from Tibetan pig MAG098^36^; Ar-2 with Chinese roe deer RGIG3983^41^ strain; and Ar-3 with Tibetan pig MAG196^36^. Likewise, we observed Ar-8 was on the same phylogenetic branch with an Australian *Methanobrevibacter gottschalkii* isolate: A27^42^(Figure 2B). Interestingly, Ar-6 formed an outbranch alongside this branch with a further outbranch containing two *Methanobrevibacter gottschalkii* strains^43, 44^. This would suggest that our resolved archaea-MAG Ar-6 might be a *Methanobrevibacter gottschalkii*. Moreover, Ar-5 clustered amongst *Methanobrevibacter smithii* strains: Tibetan pig MAG004^36^, Canadian pig SUG1019^45^, and US Florida human ATCC 35061^46^.

**Figure 1.**
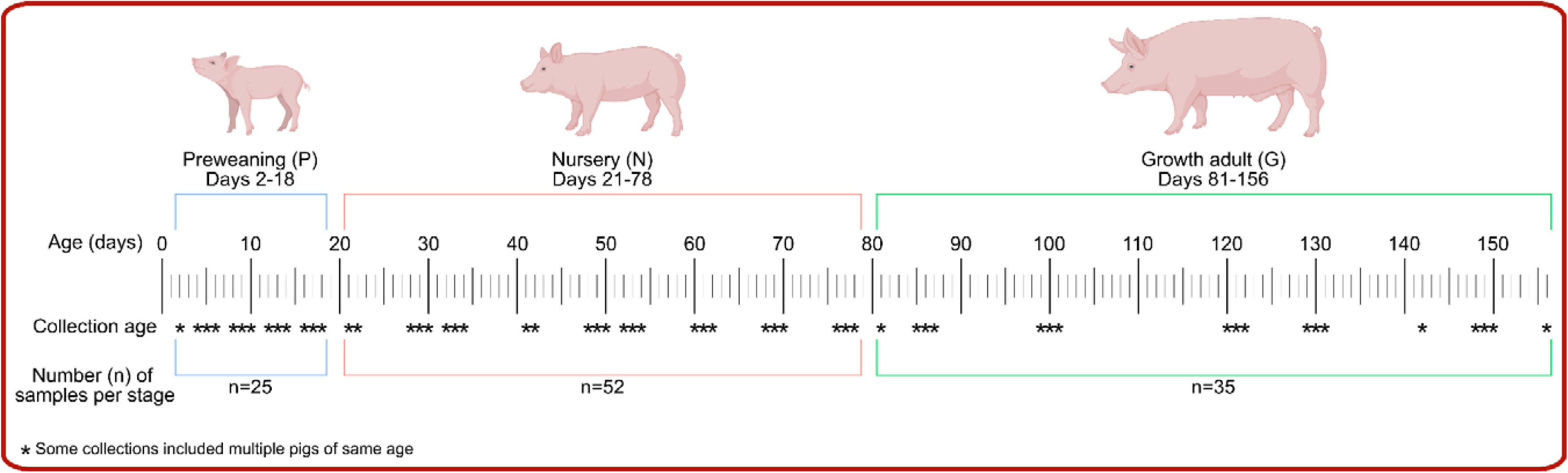
Study schematics of 7 swine hosts including fecal sampling ages and developmental st stages.

**Figure 2.**
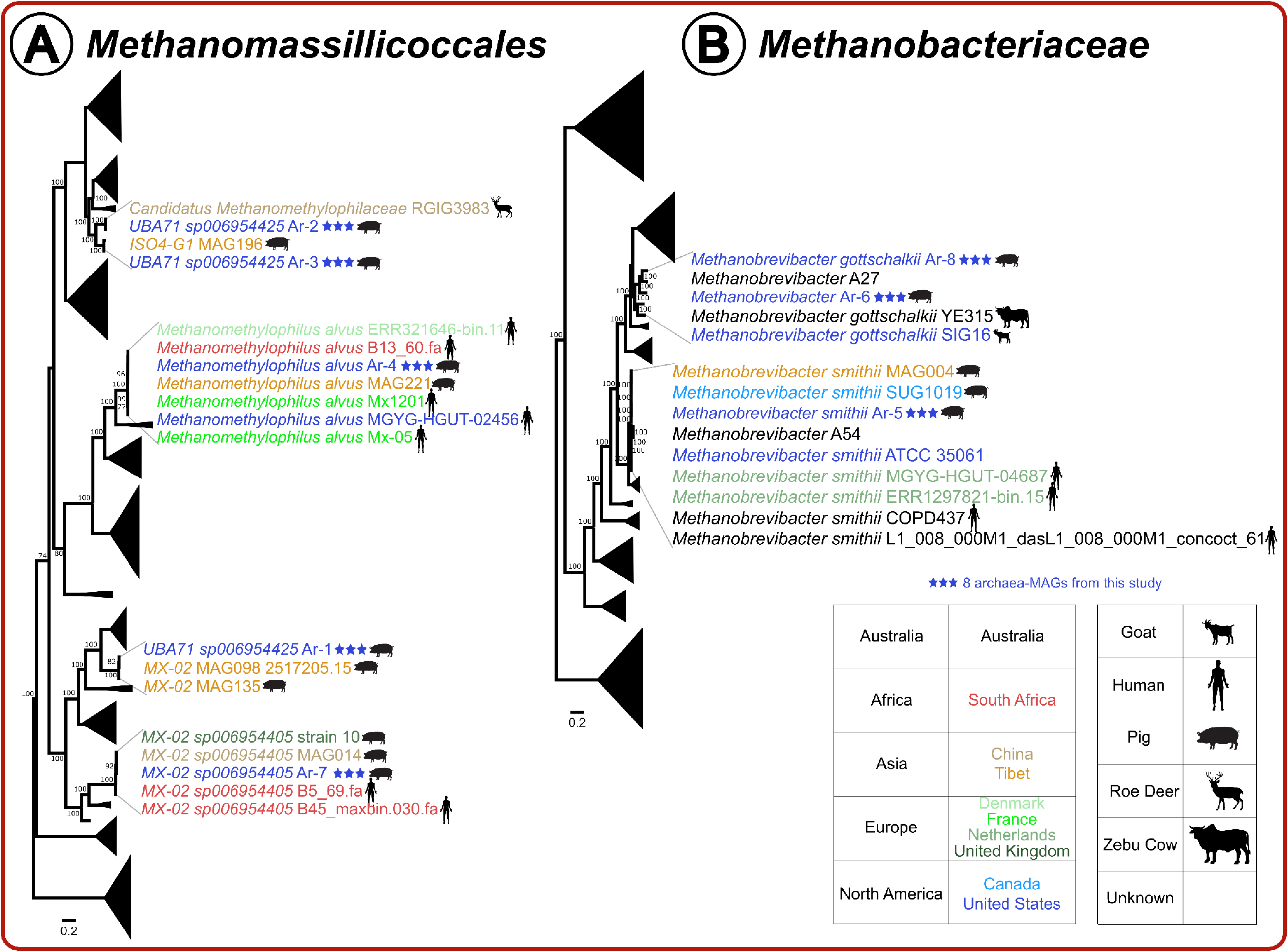
Phylogenetic trees of (A) *Methanomassiliicoccales*^33–42^ and (B) *Methanobacteriales*^36, 42–45, 126–131^ with bootstrap values ≥ 70 indicated at nodes. Branches were collapsed for non-immediate phylogenetic relatives of our archaea-MAGs while branches containing these 8 MAGs were magnified for clarity. Original trees are in Supplementary Figure S1.

While many of our methanogen MAGs clustered with swine originating archaea populations, we also demonstrated our methanogen MAGs alongside human and deer associated methanogens, suggesting similarities in microbial genes and associated functions in the methanogens amongst these host species. The roe deer similarity is especially intriguing considering that deer contain a ruminant digestive system, with four stomach compartments, compared to the single stomach system of monogastric swine and human^26, 47^. Moreover, even though our archaea-MAGs originated from the United States (in the state of Kansas) swine, our phylogenetic analyses indicated similarities to archaeal populations from Australia, South Africa, Tibet, China, United Kingdom and Canada, further supporting the global presence of archaea amongst diverse hosts. We surmise that our resolved methanogen archaea-MAGs and these close genetic relatives might be more widespread in more hosts than we expected^30^.

We recovered from our study novel archaeal genomes that were previously unidentified in US swine. We were able to resolve and obtain the genomic information, to the best of our knowledge, of the first swine-associated *Methanobrevibacter UBA71 sp006954425* and *Methanobrevibacter gottschalkii* MAGs. The methanogenic archaea family *Methanobacteriales* has been identified in many previous swine studies, with the majority of these studies utilizing 16S sequencing and/or real-time PCR identification^36, 39, 48–57^. Still, there is a lack of understanding of the *Methanobacteriales* in terms of genomic studies, and the *Methanomassiliicoccales* order collectively in general. Up to this moment, only three swine *Methanomassiliicoccales* MAGs (*Methanomethylophilus alvus*, *MX-02 sp006954405*, and *Methanobrevibacter smithii*) have been identified^36, 39, 57^. Thus, adding our highly resolved novel archaea-MAGs to the repertoire of swine-associated microbial populations will aid in understanding swine archaea, including functions, host associations (such as age, health status, sex, etc.), and global distribution.

### Prevalence of archaeal MAGs and variants at distinct host ages

Assessing the abundance of the methanogens in different growth stages of the swine host provided an opportunity to investigate the association between the host-associated methanogens and the different conditions faced by the swine as they grow. Our genome-centric metagenome analyses revealed two dominant orders of archaea - *Methanobacteriales* and *Methanomassiliicoccales*. We showed that resolved methanogen MAGs were differentially detected at different growth stages of the swine, but does the environment affect the functional niche specificity between these two orders of archaea?

The heatmap shown in Figure 3A provides a graphical summary of the changes in detection for the archaea-MAGs. Hierarchical clustering grouped the archaea-MAGs into three clusters based on detection: A (top cluster; Ar-1 through Ar-4), B (middle cluster; Ar-5 and Ar-6), and C (bottom cluster; Ar-7 and Ar-8). We observed that Cluster A contained only *Methanomassiliicoccales* MAGs, while Cluster B contained 2 *Methanobacteriales* MAGs, and Cluster C one of each order. Cluster A archaea-MAGs were primarily identified in the final stage of growth adult hosts. Conversely, Cluster B methanogens were primarily identified in preweaning hosts. Finally, Cluster C archaea were identified throughout the host lifetime.

**Figure 3.**
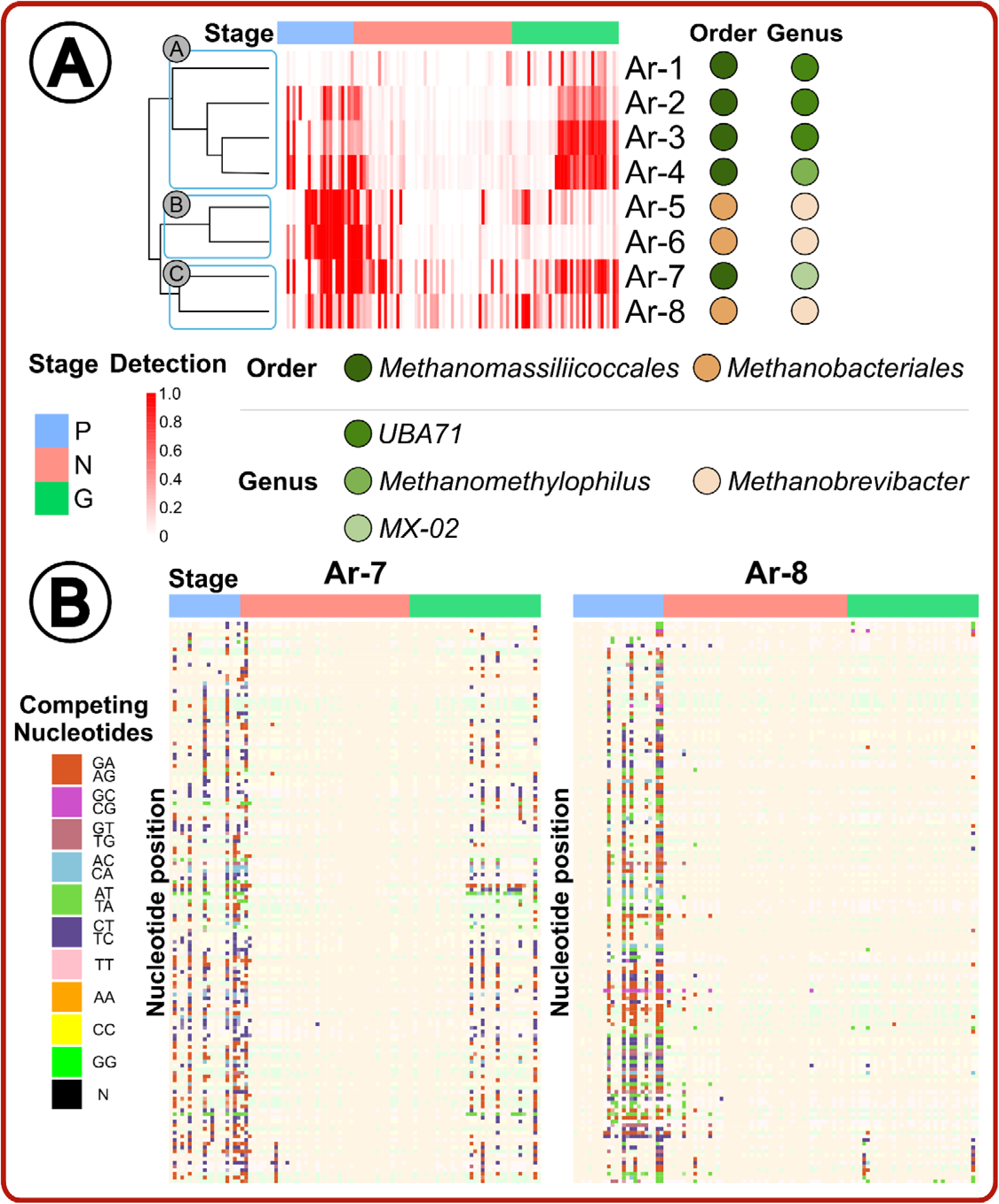
(A) Detection heatmap of archaea-MAGs (rows) across all individual sample metagenomes (columns) with MAG taxonomy and stage annotation (Preweaning [P]; nursery [N]; growth adult [G]). (B) Single-nucleotide variant (SNV) analysis of Ar-7 and Ar-8 where box colors indicate competing nucleotides and stage is indicated along the bottom.

We noticed the majority of archaea-MAGs detection values increased closely after a stage transition (preweaning to nursery and nursery to growth adult), suggesting that stage transition changes, including diet, housing, and stress, can lead to changes in microbiome composition^58^. Although, exactly how these changes impact archaea is relatively understudied, as most research evaluates bacteria, and therefore archaea- stage dynamics are a topic for future research^59, 60^.

We investigated methanogen variants, and found the majority of variation occurred in periods when other archaea were dominating (preweaning and growth adult; Supplementary Table S5). We performed single-nucleotide variant (SNV) analysis on our two archaea-MAGs that showed continuous detection throughout the host lifetime (Cluster C MAGs: Ar-7 and Ar-8; Figure 3B). We attributed the majority of variances to the unweaned host, while fewer variances were identified in the growth adult. Interestingly, the variants were highest during times where other archaea-MAGs were predominantly identified (Figure 3B). We hypothesized that the variation found in the growth adult host could indicate a competitive microbial environment, while fewer variants as compared to the earlier growth stages could be due to a largely already developed gut microbiome. In a competitive gut microbiome system, it is beneficial to have genetic diversity which translates to increased functional diversity^61^. A similar competitive environment and SNV diversity was demonstrated in the human gut bacterial community^61^. Comparatively, as the gut developed and microbes established with focused functions, the variation decreased when humans reached 2 years of age^61^. Human gut development is similar to the relatively faster development of the swine microbiome during preweaning and within 10 day days post weaning^62^. The conditions which encouraged the increased variation in the growth adult in our study could have been a change of diet, host stress, or other host-associated and environmental conditions^63^.

While we demonstrated differing archaea and SNV association with age, we were primarily interested in methanogen function. We hypothesized methanogenic function influenced our resolved methanogen MAGs’ ability to establish in the microbiome at different host stages through energy acquisition via host diet. Phylogenetic similarity in archaea across geography and hosts prompted an investigation into whether our archaea-MAGs were identified in other hosts of similar developmental ages, and therefore similar archaeal functions.

### Methanogens span host species, millennia, and geographic distance

We wanted to further demonstrate not only global and host distribution, but also temporal identification of our methanogens beyond genetic similarity, as illustrated in our phylogenetic analyses. We mapped metagenomic sequencing reads from young and aged hosts to our archaea-MAGs from the following hosts: swine (n=16)^64^, humans (n=429)^65, 66^, mice (n=60)^67^, chicken (n=71)^68^, and cattle (n=34)^69^ (Figure 4; Supplementary Tables S6 and S7). Our archaea-MAGs were identified in older humans and varying aged swine metagenomes, but not in the chicken, mice or cattle metagenomes. We also demonstrated evidence of our archaea-MAGs in the ancient human gut and global distribution. Altogether we determined within a host species, archaeal age-association appeared to be similar, but some archaea span multiple host species, and for millennia^30^. We hypothesized differential archaeal function may be essential to the gut microbiome of many modern and ancient monogastric hosts.

**Figure 4.**
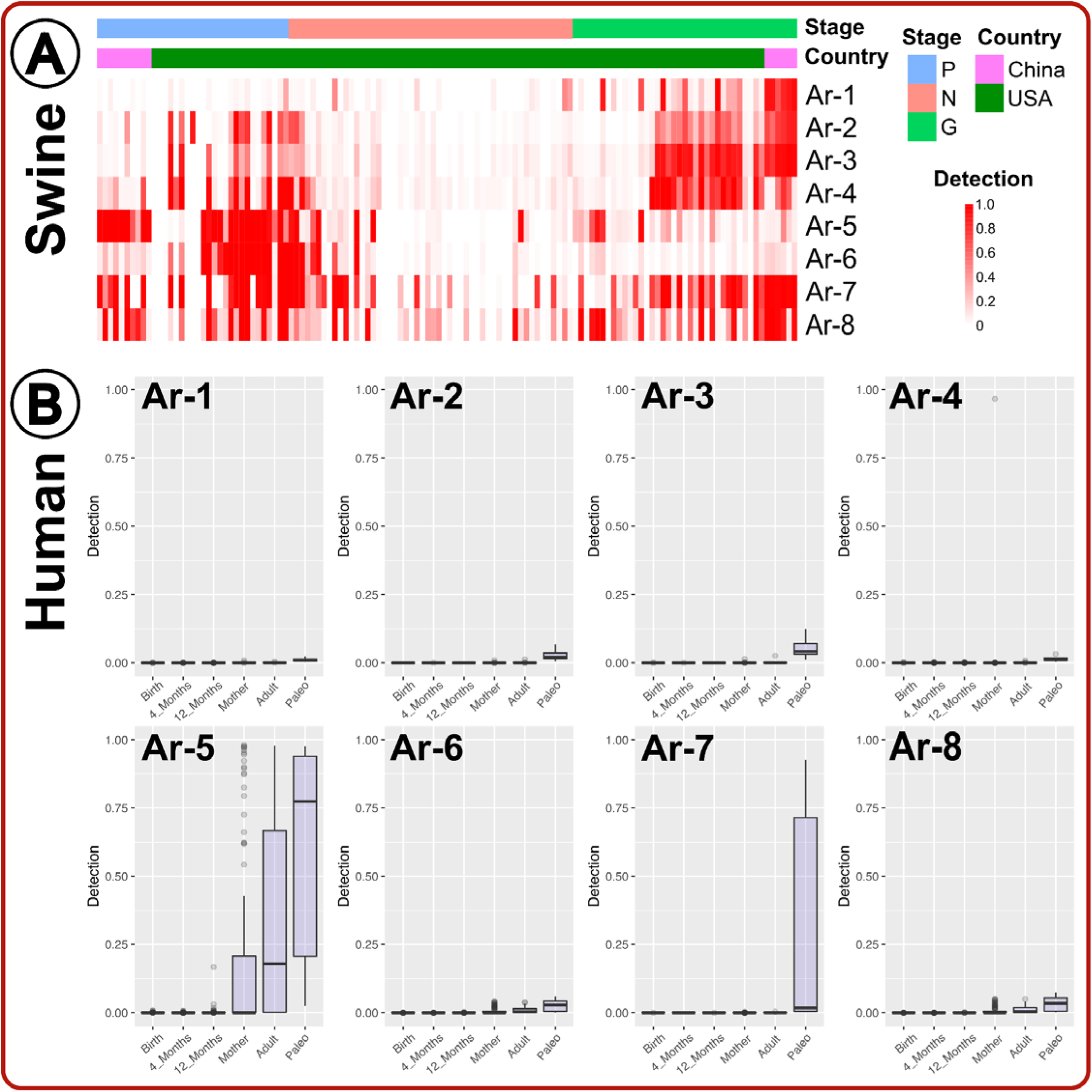
(A) Detection heatmap of previously published swine metagenomes64 mapped to this publication’s archaeal MAGs (Preweaning [P]; nursery [N]; growth adult [G]). (B) Detection box plots of previously published human metagenomes^65, 66^ mapped to our archaeal MAGs (“Adult” from Mexican humans; “Paleo” from present day US and Mexico ; all remaining groups from Sweden).

We determined our swine-associated methanogens were not present in poultry, mice and ruminant metagenomes (Supplementary Table S7). Given the drastic differences in the ruminant digestive system compared to the monogastric gut, we were not surprised that our swine archaea-MAGs were not found in cattle from the United States (US) State of Pennsylvania. Although not identified consistently in all cattle, *Methanobrevibacter smithii*^70–73^ and *Methanobrevibacter gottschalkii*^74–77^ have been associated with the cow digestive tract. Similarly, *UBA71* has been identified in adult chickens previously^78^. Given that similar taxonomic methanogens are present in cattle and chickens, we hypothesized the methanogens of these hosts were genetically distinct from the methanogens we identified in swine. Additionally, since the methanogens we identified were not consistently detected across our aging hosts, it was very probable that other metagenomes from these host populations could contain our methanogens. Future research is necessary to evaluate how distinct methanogen members function individually and collectively within the microbiome system to influence gut health in different host species.

Interestingly, we could only find a singular example of archaea attributed to the mouse gut: *Methanomassiliicoccaceae DTU008*^79^. Remaining attempts, encompassing more than 1,000 metagenomes, proved unsuccessful in identifying mice gut archaea^80–83^. In fact, an investigation of murine gut composition across 17 rodent species demonstrated, beyond the instance of mouse *DTU008*, only North American porcupine (*Erethizon dorsatum*), capybara (*Hydrochoerus hydrochaeris*), and guinea pig (*Cavia porcellus*) contained archaea^79^. Although murine hosts have a monogastric digestive system, there appears to be a lack of understanding if and when archaea are present in mouse gut^83^.

Although from a different continent, Chinese swine demonstrated the closest age-associated detection to our US swine methanogens. Even though the Chinese preweaning swine were not weaned, the methanogen presence appeared to more closely resemble the US swine weaned, nursery gut. The exception to the nursery resemblance being Ar-5, which more closely resembled our US swine preweaning gut. Many factors, including breed, weaning age (China at 42 days; US at 18-20 days), and housing, are known to influence microbiome development, and therefore could have resulted in the different archaeal age-establishment patterns^58, 84, 85^. In terms of our earlier detection clusters, Cluster A still appeared more prevalent in the growth adult host, and Cluster C was similarly prevalent throughout both the preweaning and growth adult stages. Although the detections of Ar-5 and Ar-6 (Cluster B) were not shared between the Chinese and US swine, as Ar-6 demonstrated relatively low detection in the preweaning stage. Given that the Chinese swine dataset was from a single day in preweaning and growth adult, future research should investigate longitudinal distribution of methanogens from monogastric swine according to various characteristics, such as country of origin, breed, diet, housing environment, etc. This would further develop our understanding of global methanogen distribution according to associated variables.

Overall, we demonstrated genetic support for the same, or very closely related, methanogens circulating in both US and Chinese swine with similar age-associated detection. This further demonstrates the ubiquity of archaea to the monogastric swine host which we hypothesized are distributed with host age according to archaeal function.

Our resolved archaea-MAGs not only appeared in Chinese swine, but we also provided evidence of these archaea-MAGs in adult humans from modern age (Mexico and Sweden) and ancient time (modern day US and Mexico). In contrast to the swine gut, we identified merely two methanogens in the human gut: *M. smithii* and *MX-02 sp006954405*. With the exception of one Swedish 12 month sample demonstrating Ar-5 presence, the infant data illustrated comparatively minimal to no archaeal presence of our methanogens. Given that many publications demonstrate identification of multiple *Methanobacteriales* and *Methanomassiliicoccales* in the human gut^4, 6, 86, 87^, it is possible that there were genetically distinct methanogens present in these human samples beyond our *M. smithii* and *MX-02 sp006954405*. The modern adult human samples, both the Mexican and Swedish datasets, only demonstrated *M. smithii* presence. Multiple publications have identified increasing *M. smithii* in the human gut with age^87, 88^. Interestingly, we identified *MX-02 sp006954405* and *M. smithii* in the palaeofaeces from the US and Mexico, suspected to be between 1,000-2,000 years old^65^. While many paleobiology studies have investigated ancient methanogens of water sediments^89–97^, we identified merely two human-related paleobiology methanogen studies pertaining to: the archaic human gut^65^ and neanderthal dental plaques^98^. *M. smithii* has been previously identified in ancient humans^65^, but the identification of *MX-02 sp006954405* appeared to be the first evidence of human-associated ancient *Methanomassiliicoccales*. The methanogen *MX-02* has been identified in the human gut previously^18^, but we illustrated novel evidence for *MX-02 sp006954405* in the ancient human gut, suggesting that *MX-02 sp006954405*, or close relatives, were likely present in the modernized human gut, but we did not identify genetic resemblance in the 122 modern human metagenomes we evaluated. Future research is necessary to provide further insights into the gut-associated archaea to elucidate genetic phylogeny, evolution of archaeal functions, and association with ancient humans.

Given our findings indicating our US-swine associated archaea-MAGs were present in Chinese swine, US humans, and Mexican humans, we wanted to further understand the role of these methanogens in the monogastric gut.

### Critical methane metabolism functions were conserved across methanogens

Our primary goal was to profile the expressed genomic potential that contributed to the methanogenesis of the swine gut. In our study, we constructed and demonstrated the first swine complete methanogenic pathways, which is crucial for understanding the role of archaea within the microbiome system and to host health. We analyzed methanogenesis pathways of *Methanobacteriales* and *Methanomassiliicoccales* (Figure 5, Supplementary Table S4). We identified 44 genes in methane metabolism^99^.

**Figure 5.**
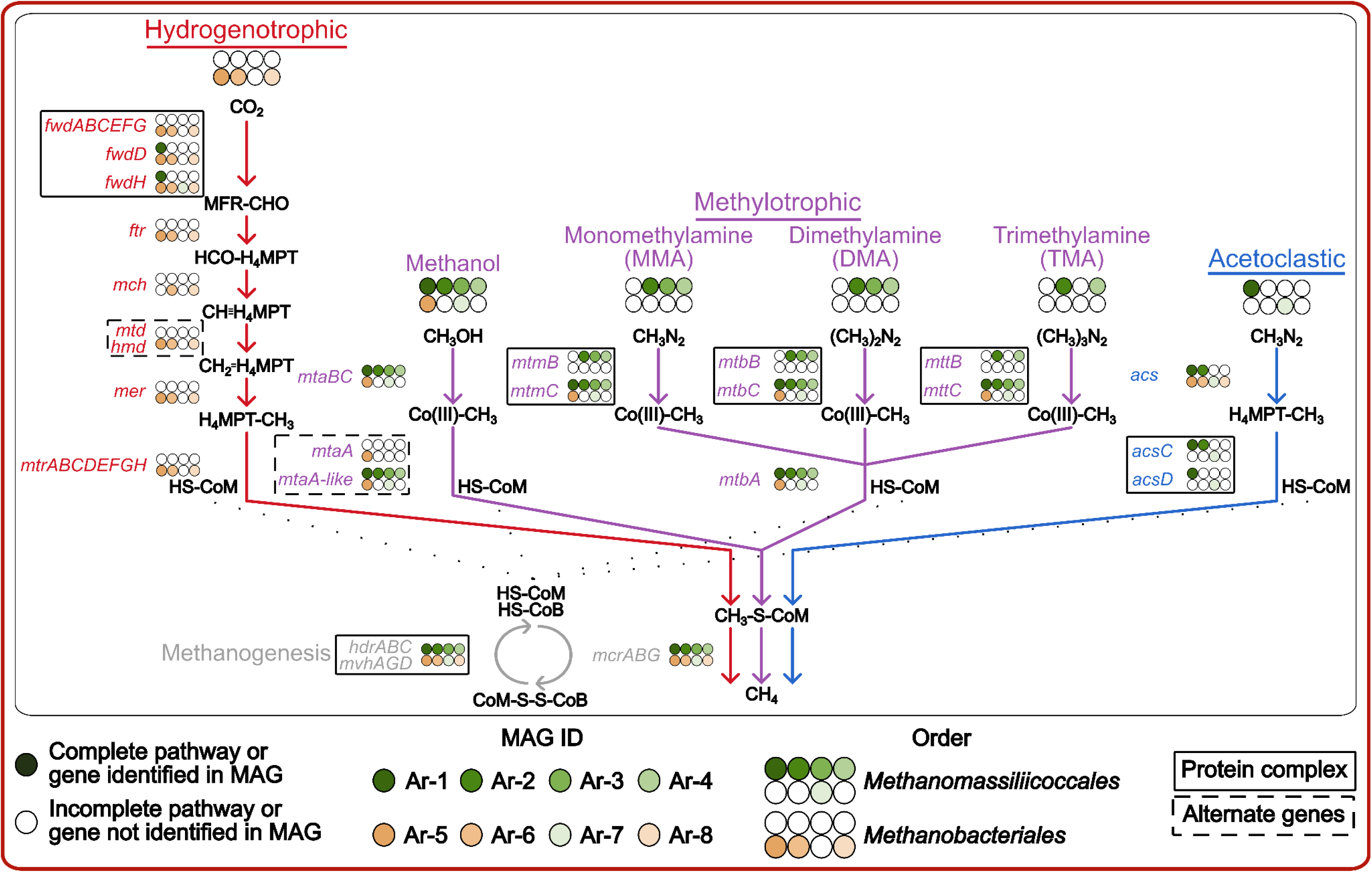
Methane metabolic pathway genes detected in our archaeal MAGs distinguished by pathway^32, 111, 132, 133^.

The shared genes represented crucial functions in the final methanogenesis steps of energy and methane production. Nine genes were shared across the 8 archaea-MAGs: 3 heterodisulfide reductase (*hdrA*, *hdrB*, and *hdrC*), 3 methylviologen-dependent Ni,Fe hydrogenase (*mvhA*, *mvhG*, and *mvhD*), and 3 methyl-coenzyme M reductase (*mcrA*, *mcrB*, and *mcrG*). HdrABC, MvhAGD, and McrABG are critical to the final steps of methanogenesis. HdrABC and MvhAGD form an electron-bifurcating complex to regenerate coenzyme M (HS-CoM) and coenzyme B (HS-CoB) from heterodisulfide CoM-S-S-CoB^100^. An intermediate step, not shared by all archaea-MAGs but discussed in the following section, generates methylated cozyme M (CH_3_-S-CoM)^27^. McrABG then catalyzes the final methanogenesis step where the methyl group of CoM-CH_3_ is reduced to methane, with HS-CoB utilized as an electron donor^27^. This formation of methane generates CoM-S-S-CoB for reduction again by HdrABC/MvhAGD^101^. The final step in producing methane is crucial for methanogens, and our archaea-MAGs supported the roles of HdrABC, MvhAGD, and McrABC in these final methanogenesis reactions.

### Differential Methanobacteriales and Methanomassiliicoccales methane metabolic pathways may relate to age-associated detection

While we supported the clear ubiquity of *hdr*, *mvh* and *mcr* to methanogens, we were primarily interested in how our archaea differed in methanogenic potential since they exhibited differential colonization through the swine growth stages. We noticed our taxonomically distinct archaea-MAGs harbored different genetic components for divergent methane metabolic pathways (Figure 5). We identified the first genetic support of an acetoclastic *Methanomassiliicoccales*, and first acetoclastic methanogen in a monogastric host. Moreover, *Methanomassiliicoccales* also contained genes for the methylotrophic methanogenic pathway, indicating the potential to utilize various substrates from acetate to methylated compounds. Alternatively, *Methanobacteriales* contained genes for the hydrogenotrophic pathway with CO_2_ as the substrate input. We surmised that these alternate pathways played a role in energy acquisition according to differential nutrients available during the host lifetime, and therefore growth stage associated diet.

The hydrogenotrophic methanogenic pathway was the key methanogenic pathway identified in all of our *Methanobacteriales* MAGs. During hydrogenotrophic methanogenesis, CO_2_ is reduced to methane (CH_4_) with four molecules of H_2_ . Our *Methanobacteriales* archaea-MAGs contained all genes crucial to the hydrogenotrophic methanogenic pathway: formylmethanofuran dehydrogenase (*fwd*; A-H), formylmethanofuran-tetrahydromethanopterin N-formyltransferase (*ftr*), methenyltetrahydromethanopterin cyclohydrolase (*mch*), coenzyme F_420_-dependent methylenetetrahydromethanopterin dehydrogenase (*mtd*), methenyltetrahydromethanopterin hydrogenase (*hmd*), methylenetetrahydromethanopterin reductase (*mer*), and methyltetrahydromethanopterin-coenzyme M methyltransferase (*mtr*; A-H)^32, 103^.

We identified both *mtd* and *hmd*, indicating the methanogens can potentially utilize H_2_ with or without F_420_ to reduce methenyl-H_4_MPT, as Mtd requires F_420_^104^. As mentioned previously, we believe the CH_3_-CoM in our *Methanobacteriales* archaea-MAGs was generated via MtrABCDEFGH. This *mtr* complex has been demonstrated to transfer the methyl group from tetrahydromethanopterin (H_4_MPT) to CoM, therefore coupling the hydrogenotrophic pathway to the final methane production steps^14, 105, 106^. *Methanobacteriales* have been associated with the hydrogenotrophic pathway previously^27^.

Our *Methanobacteriales* archaea-MAGs contained all of the aforementioned genes, with one exception: Ar-5 lacked *mch*. We attribute this to the incompleteness of the metagenome assembled genome, as Ar-5 exhibited the lowest completion of our methanogen MAGs at ∼72% (Table 1, Supplementary Table S3). Interestingly, only two hydrogenotrophic pathway genes (*fwdD* and *fwdH*) were identified in two *Methanomassiliicoccales* archaea-MAGs. The lack of a hydrogenotrophic pathway clearly indicated the *Methanomassiliicoccales* utilized a distinct methane pathway.

All *Methanomassiliicoccales* archaea-MAGs, and our *M. smithii* archaea-MAGs, indicated varying ability to metabolize methanol, mono-, di- and trimethylamine through the methylotrophic methanogenesis pathway. Genes we identified in our archaea-MAGs included: methanol:coenzyme M methyltransferase (*mtaB* and *mtaC*), monomethylamine (MMA) methyltransferase (*mtmB* and *mtmC*), dimethylamine (DMA) methyltransferase (*mtbA*, *mtbB*, and *mtbC*), and trimethylamine (TMA) methyltransferase (*mttB* and *mttC*)^32^. *Methanomassiliicoccales* is known to perform methylotrophic methanogenesis with all of the previously discussed substrates^107^. Conversely, *M. smithii* was thought to be a hydrogenotrophic population^108^. Given we only identified methylotrophic genes in one *Methanobacteriales* population, future research is necessary to support this genetic potential.

Although all *Methanomassiliicoccales* methanogens and the *M. smithii* population contained the *mtaBC* genes for methanol metabolism, we did not identify *mtaA* in the *Methanomassiliicoccales* genetic content. The MtaABC complex transfers the methyl group from methanol to coenzyme M, generating CH_3_-S-CoM for McrABG reduction^109^. We identified a candidate *mtaA* homolog through a literature review: uroporphyrinogen III decarboxylase (*hemE*)^110^ (designated *mta-like* in Figure 5 and Supplementary Table 4). This is the first publication identifying the homolog in methanogens. As such, future research is critical to analyze how HemE might interact with MtaBC, and how the enzyme performs in the methylotrophic pathway.

Only three (Ar-2, Ar-3, and Ar-4) out of the five *Methanomassiliicoccales* archaea-MAGs contained a complete genetic pathway associated with monomethylamine (*mtmBC*) and dimethylamine (*mtbBC*) utilization^32, 111^. Moreover, only two *Methanomassiliicoccales* populations (Ar-2 and Ar-4) contained *mttBC*, associated with trimethylamine (TMA) utilization^32, 111^. TMA has been associated with increased cardiovascular disease, so TMA metabolism is beneficial for the host^4, 5^. Further research is necessary to evaluate if other methylated compounds may play roles in cardiovascular disease, and therefore the host aided by archaeal metabolism. Although the majority of our archaea-MAGs appeared to be able to utilize methanol in the methylotrophic methanogenesis pathway, fewer were able to use mono-, di- and trimethylamine. As noted previously, this might be due to the incompleteness of the archaea-MAGs, especially since our *Methanomassiliicoccales* archaea-MAGs completeness ranged from ∼80-99% (Table 1, Supplementary Table S3). Still, there is a possibility that these *Methanomassiliicoccales* archaea-MAGs might have different abilities in utilizing methylated sources due to contrasting biological necessity and evolutionary selection^112^.

To the best of our knowledge, we identified the first complete acetoclastic methanogenic pathway in *Methanomassiliicoccales*^113, 114^. Three *Methanomassiliicoccales* (Ar-1, Ar-2, and Ar-7) archaea-MAGs contained genetic support, with two complete pathways (Ar-1 and Ar-7), for the acetoclastic, or also called aceticlastic, pathway. Acetate is reduced to acetyl-CoA and then H_4_MPT via acetyl-CoA synthetase (*acs*) and carbon monoxide dehydrogenase (*acsC* and *acsD*)^32, 111, 115^. Two of our *Methanomassiliicoccales* archaea- MAGs (Ar-3 and Ar-4) did not have acetoclastic genes identified, while *acsD* was not found in Ar-2.

Acetoclastic methanogenesis is typically performed by aquatic methanogens^116–118^. The only prior identification of acetoclastic archaea in gastrointestinal tracts was in *Methanosarcinales* of a ruminant (cow)^4, 119^. Therefore, we identified the first evidence for acetoclastic methanogenesis in monogastrics. Moreover, only *Methanosarcinales* and *Methanococcales* were known to perform acetoclastic methanogenesis^113, 119, 120^. The ability of our *Methanomassiliicoccales* to perform both methylotrophic and acetoclastic methanogenesis parallels *Methanosarcinales*, since many *Methanosarcinales* also are able to perform both of these pathways^114^. Conversely, acetoclastic-able *Methanococcales* are also able to perform hydrogenotrophic methanogenesis^119^. Acetoclastic methanogenesis requires an ATP input to convert acetate to acetyl-CoA, which impairs the energy efficiency of this methanogenic pathway^121^. The diminished energy return of acetoclastic methanogenesis likely plays a role in the dual acetoclastic-methylotrophic metabolic potential of our *Methanococcales* archaea-MAGs. The ability to utilize different substrates via varying methanogenic pathways is beneficial. The methanogen can potentially still metabolize energy as their substrate source changes. Changing substrates is common in the gastrointestinal system, as the host changes diets and the associated microbiome changes and produces different metabolites^59^.

Although the *Methanomassiliicoccales* archaea-MAGs contained genes for both methylotrophic and acetoclastic methanogenesis, this dynamic substrate capability did not appear to allow *Methanomassiliicoccales* archaea-MAGs to prevail more than their counterparts: *Methanobacteriales*. In fact, the *Methanobacteriales* populations in general were detected at more times during the host life than *Methanomassiliicoccales* (Figure 3A). This may indicate CO_2_ is more available in the monogastric system during the host lifetime. The exception to this being the during the growth stage where *Methanomassiliicoccales* appear abundantly, therefore mylated compounds or acetate could have transitioned to being the dominantly available substrate, suggesting that the stage-associated characteristics, such as dietary composition had a high influence on the methanogens dynamics. Although diet influences substrate availability, we cannot rule out other factors, including host age and other gastrointestinal organisms (including bacteria and protist), which alter the gut microbiome system^4, 59, 119, 122^.

Collectively, with the novel acetoclastic *Methanomassiliicoccales* and methylotrophic *Methanobacteriales M. smithii* archaea-MAGs, there is a knowledge gap surrounding the functional potential of methanogens. Taken together with the phylogenetic analysis, we are lacking a holistic understanding from global and host distribution of methanogens to their methanogenic actions. Other publications have also discussed the lack of overall methanogen knowledge, especially knowledge surrounding archaeal functions^123–125^. Archaea are a member of the gut microbiome alongside bacteria, fungi and viruses, and without understanding their distribution and functions, we will not understand how archaea influence the gut microbiome system and host health.

## Conclusions

We performed a longitudinal study of the monogastric microbiome where we produced 1,130 MAGs, with 8 methanogen archaea-MAGs. The novel archaea-MAGs clustered with geographically diverse methanogens from various animal and human hosts, indicating global distribution of closely related archaea. We also determined that our archaea-MAGs were detected in swine and humans from distinct continents and time. Given the stark distinctions in detection and distribution, we wanted to evaluate if energy acquisition associated with methanogenesis could be related to these factors, especially age-distribution. Our *Methanobacteriales* archaea-MAGs contained genes for hydrogenotrophic methanogenesis, indicating the ability to metabolize CO_2_. Alternatively, *Methanomassiliicoccales* archaea-MAGs appeared to have the capability to utilize a range of substrates from methylated compounds, including methanol and methylamine, and acetate, through the methylotrophic and acetoclastic pathways, respectively. We identified the first acetoclastic *Methanomassiliicoccales*, and also the first acetoclastic methanogens of monogastrics. Moreover, we identified a *Methanobacteriales* population with methylotrophic genes. Previously, *Methanobacteriales* was thought to only perform hydrogenotrophic methanogenesis. We hypothesized the distinct diets, given according to age, provided different substrates which influenced archaeal establishment and therefore detection patterns. Still, we know there are multiple other growth stage-associated and microbiome dynamics which likely play a role in archaeal growth.

In order to continue developing our understanding of archaea, we must continue to evaluate their global prevalence across diverse hosts and ecosystems. Moreover, we should evaluate the significance of acetoclastic methanogens to monogastrics, including how these methanogens influence other microorganisms and host health. Future studies should also investigate how growth stage-associated factors influence methanogenic potential and therefore archaeal abundance. In pursuing this archaea research, we can better determine how methanogens provide beneficial or detrimental consequences to host health, and how we might utilize or deter methanogens in animals and humans alike.

## Supplementary Files

Supplementary Table S1. Demographics (diet, birth date, housing group, etc.) of swine hosts and dams, and sample metadata (swine age, host ID, stage and general health information, etc.).

Supplementary Table S2: Raw sequencing analysis by QIIME2^134^, including 16S rRNA counts initially obtained, counts after primer trimming, DADA2 quality control per sample, and NCBI sample accession.

Supplementary Table S3. Anvi’o results from initial bins and resulting redundant and nonredundant MAGs, including taxonomic classification detailing archaeal MAGs and PATRIC genome IDs^135^.

Supplementary Table S4. PATRIC RAStk and COGG gene annotations for methanogen MAGs.

Supplementary Table S5. Single nucleotide variant (SNV) results.

Supplementary Table S6. Metadata from metagenomes utilized in mapping to our archaeal MAGs (swine^64^, human infant^66^, human adult and palaeofaeces^65^, mice^67^, chicken^68^, and cattle^69^).

Supplementary Table S7. Detection results from metagenome mapping to archaea- MAGs.

Supplementary Figure S1. Complete, non-collapsed, *Methanobacteriales* and *Methanomassiliicoccales* phylogenetic trees with statistics.

## Materials and Methods

### Study design, sample collection and DNA extraction

Our study design and sample collection occurred as previously described^136^. We collected fecal samples from 7 swine over 22 timepoints, ranging in swine age from 1 to 156 days across three developmental stages: preweaning (P), nursery (N), and growth adult (G) (Figure 1, Supplementary Table S1). Swine were born and raised at the Kansas State University Swine Teaching and Research Center. Swine originated from the same farrowing group, and were weaned between 18-20 days of age, depending on day of birth. Pigs were managed according to the Kansas State University Institutional Animal Care and Use Committee (IACUC) approved protocol #4036, and methods are reported according to ARRIVE guidelines. The authors also confirmed that all methods were performed in accordance with relevant guidelines and regulations, and we affirmed that all methods were approved by Kansas State University.

We stored fecal samples at -80°C until DNA extraction. We extracted total genomic DNA from fecal samples utilizing the E.Z.N.A.® Stool DNA Kit (Omega Bio-tek Inc.; Norcross, GA), following the manufacturer protocols. We then quantified the extracted genomic DNA with a Nanodrop and Qubit™ (dsDNA BR Assay Kit [Thermo Fisher; Waltham, MA]) for DNA quality and concentration. We stored extracted DNA at -80°C until library preparation and sequencing.

### Metagenomic sequencing and ‘omics workflow

DNA libraries were generated for a total of 112 samples with Nextera DNA Flex (Illumina, Inc.; San Diego, CA). Resulting libraries were then visualized on a Tapestation 4200 (Agilent; Santa Clara, CA) and size-selected using the BluePippin (Sage Science; Beverly, MA). The final library pool of 112 samples was quantified on the Kapa Biosystems (Roche Sequencing; Pleasanton, CA) qPCR protocol, and sequenced on the Illumina NovaSeq S1 chip (Illumina, Inc.; San Diego, CA) with a 2 x 150 bp paired-end sequencing strategy.

We utilized the ‘anvi-run-workflow’ program to run a combined bioinformatics workflow in anvi’o v.7.1 (https://anvio.org/install/)^135,137^, with a co-assembling strategy. The workflow used Snakemake to implement numerous tasks including: short-read quality filtering, assembly, gene calling, functional annotation, hidden Markov model search, metagenomic read-recruitment and binning^138^. Briefly, we processed sequencing reads using anvi’o’s ‘iu-filer-quality-minoche’ program, which removed low-quality reads following criteria outlined in Minoche *et al*.^139^. The resulting quality-control reads were termed “metagenome” per sample. We organized the samples into 3 metagenomic groups based on the developmental stages (P, N, G), and used anvi’o’s MEGAHIT v1.2.9 to co-assemble quality-filtered short reads into longer contiguous sequences (contigs)^135, 140^. The following methods were then utilized in anvi’o to further process the contigs: (1) ‘anvi-gen-contigs-database’ to compute k-mer frequencies and identify open reading frames (ORFs) using Prodigal v2.6.3^135, 141^; (2) ‘anvi-run-hmms’ to annotate bacterial and archaeal single-copy, core genes using HMMER v.3.2.1^135, 142^; (3) ‘anvi-run-ncbi-cogs’ to annotate ORFs with NCBI’s Clusters of Orthologous Groups (COGs; https://www.ncbi.nlm.nih.gov/research/cog)^143^; and (4) ‘anvi-run-kegg-kofams’ to annotate ORFs from KOfam HMM databases of KEGG orthologs (https://www.genome.jp/kegg/)^144^.

We mapped metagenomic short reads to contigs in anvi’o with Bowtie2 v2.3.5^145^, and we then converted mappings to BAM files with samtools v1.9^135, 146, 147^. We used the anvi’o ‘anvi-profile’ program to profile BAM files with a minimum contig length of 1,000 bp. Next, we combined profiles with ‘anvi-merge’ into a single anvi’o profile for downstream analyses. We grouped contigs into bins with ‘anvi-cluster-contigs’ and CONCOCT v1.1.0^148^. We manually processed bins with ‘anvi-refine’ using bin tetranucleotide frequency and coverage across samples^135, 149, 150^. Following manual processing, we labeled bins that had >70% completion and <10% redundancy (both based on single-copy core gene annotation) as metagenome-assembled genomes (MAGs). Finally, we used ‘anvi-compute-genome-similarity’ to calculate average nucleotide identity (ANI), using PyANI v0.2.9^135, 151^, for each MAG to identify non-redundant MAGs. We analyzed MAG occurence in a sample with the “detection” metric.

We considered a MAG as detected in a metagenome if the detection was >0.25, which is an appropriate cutoff to eliminate false-positive signals in read recruitment results. We used ‘anvi-gen-variability-profile’ with ‘--quince-mode’ to export single-nucleotide variant (SNV) information on all MAGs after read recruitment, to identify subpopulations of the MAGs in the metagenomes^135^. We used DESMAN v2.1.1 in anvi’o to analyze the SNVs and determine the number and distribution of subpopulations in the MAGs^152^. We accounted for non-specific mapping by removing any subpopulations that made up less than 1% of the entire population that were explained by a single MAG.

### Data analyses

We used the “detection” criteria (>0.25) for downstream statistical analyses. We downloaded metagenomes from swine^64^, humans^65, 66^, mice^67^, chicken^68^, and cattle^69^, and performed mapping to the non-redundant archaea-MAGs according to specifications above (Supplementary Table S2). We used RStudio v1.3.1093^153^ to visualize MAGs detection patterns in RStudio (https://www.rstudio.com/products/rstudio/) using: pheatmap (pretty heatmaps) v1.0.12^154^, ggplot2 v3.3.5 (https://ggplot2.tidyverse.org/)^155^, forcats v0.5.1 (https://forcats.tidyverse.org/)^156^, dplyr v1.0.8 (https://dplyr.tidyverse.org/)^157^, and ggpubr v0.4.0 (https://CRAN.R-project.org/package=ggpubr)^158^.

We utilized the RASTtk Genome Annotation Service on PATRIC v3.6.12 (https://patricbrc.org/) and anvi’o COG annotations for metabolic function analyses^159, 160^. We used the comparative pathway tool in PATRIC to predict the metabolic pathways of our resolved non-redundant MAGs. We obtained similar genomes that were deposited in public databases and performed phylogenetic analyses of our non-redundant MAGs in PATRIC^160^. Parameters were set as follows: 100 genes, 10 max allowed deletions, and 10 max allowed duplications. We constructed phylogenetic trees for our MAGs with 192 closely related genomes, using the amino acid and nucleotide sequences from the global protein families database. RAxML program was used to construct the trees based on pairwise differences between the aligned protein families of the selected sequences.

Our final figures were edited in Inkscape v1.2.1^161^.

### Data availability

We uploaded our metagenome raw sequencing data to the SRA under NCBI BioProject PRJNA798835. All other analyzed data, in the form of databases and fasta files, and bioinformatic scripts are accessible at figshare 10.6084/m9.figshare.20431713.

## Acknowledgements

Our team is very grateful to the large number of individuals and organizations which assisted us in performing this research. We thank members of the Kansas State University swine team (Frank Martin, Mark Nelson, Duane Baughman, and Julia Holen) for aiding in the sample collection. Gratitude is also extended to the University of Kansas Medical Center Genome Sequencing Facility for their expertise and assistance in sequencing including: Clark Bloomer, Dr. Veronica Cloud, Rosanne Skinner, and Yafen Niu. We greatly appreciate assistance from the following sources: Kansas State University Interdepartmental Genetics Program (fellowship for Brandi Feehan), Global Food Systems Seed Grant Program, Kansas Intellectual and Developmental Disabilities Research Center (NIH U54 HD 090216), the Molecular Regulation of Cell Development and Differentiation – COBRE (P30 GM122731-03) - the NIH S10 High-End Instrumentation Grant (NIH S10OD021743) and the Frontiers CTSA grant (UL1TR002366) at the University of Kansas Medical Center, Kansas City, KS 66160.

## Author Contributions

B.F., R.G. and S.T.M.L. designed the study. Sample collection was performed by B.F. B.F., V.D., K.R., and K.W. completed DNA extraction and Nanodrop and Qubit quality analysis. B.F. and Q.R. performed anvi’o and PATRIC bioinformatic analyses. B.F. and S.T.M.L. attributed biological relevance, wrote the manuscript, prepared figures, and supplementary files. B.F. and S.T.M.L. performed major manuscript and figure refinement while remaining authors contributed to lighter refinement. All authors read, contributed to manuscript revision, and approved the submitted version.

## Materials & Correspondence

Requests for materials and correspondence should be addressed to Dr. Sonny T M Lee at.

