## Supplementary Figure S1 for "Novel Complete Methanogenic Pathways in Longitudinal Genomic Study of Monogastric Age-Associated Archaea"

### Phylogenetic Tree Report

#### Rendered Tree

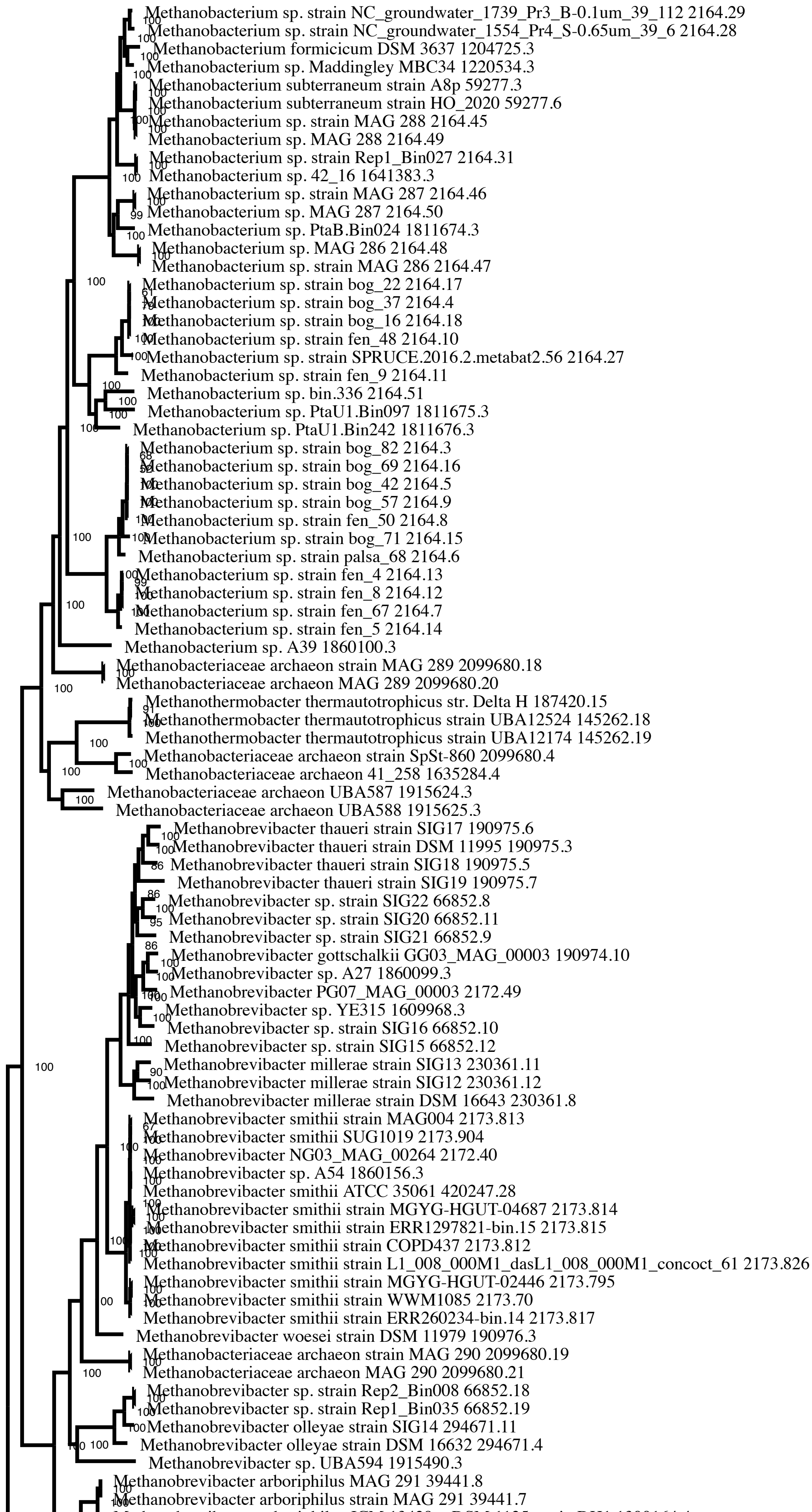

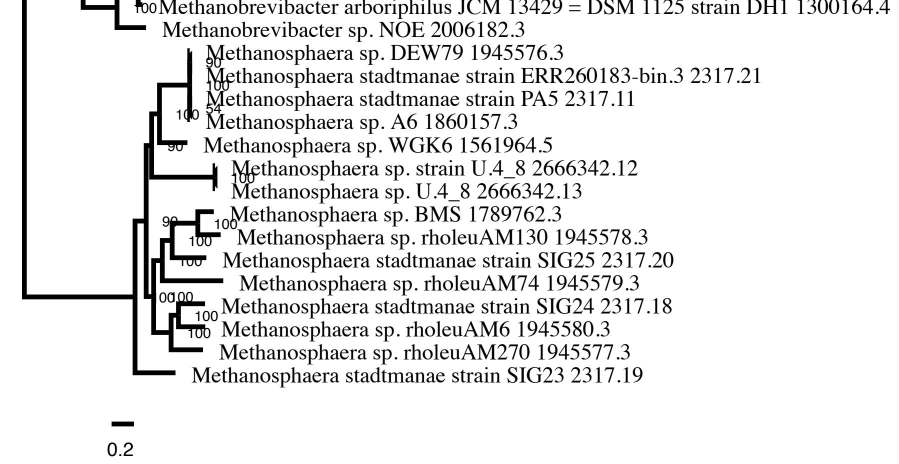


Alternate View

Files with more details

### Tree Analysis Statistics

| | |
|---|---|
| **Requested genomes** | 100 |
| **Genomes with data** | 100 |
| **Max allowed deletions** | 10 |
| **Max allowed duplications** | 10 |
| **Single-copy genes requested** | 100 |
| **Single-copy genes found** | 100 |
| **Num protein alignments** | 100 |
| **Alignment program** | mafft |
| **Protein alignment time** | 2245.4 seconds |
| **Num aligned amino acids** | 38462 |
| **Num CDS alignments** | 96 |
| **Num aligned nucleotides** | 111951 |
| **Best protein model found by RAxML** | LG |
| **Branch support method** | RAxML Fast Bootstrapping |
| **RAxML likelihood** | -4785827.9582 |
| **RAxML version** | 8.2.11 |
| **RAxML time** | 116639.0 seconds |
| **Total time** | 119167.4 seconds |

### RAxML Command Line

Goal: Analyze proteins with model 'AUTO' to find best substitution model.
raxmlHPC-PTHREADS-SSE3 -s Methanobacteriaceae_10_MAX_DELs_DUPs_proteins.phy -n Methanobacteriaceae_10_MAX_DELs_DUPs_proteins -m PROTCATAUTO -p 12345 -T 12 -e 10
Process time: 20294.419 seconds

Goal: Find best tree.
raxmlHPC-PTHREADS-SSE3 -s Methanobacteriaceae_10_MAX_DELs_DUPs.phy -n Methanobacteriaceae_10_MAX_DELs_DUPs -m GTRCAT -q Methanobacteriaceae_10_MAX_DELs_DUPs.partitions -p 12345 -T 12 -f a -x 12345 -N 100
Process time: 96344.551 seconds

### RAxML Codon and Amino Acid Partitions

```
DNA, codon1 = 1-111951\3
DNA, codon2 = 2-111951\3
DNA, codon3 = 3-111951\3
LG, proteins = 111952-150413
```

### RAxML Warnings

Sequences 2099680.18 and 2099680.20 are exactly identical
Sequences 2164.45 and 2164.49 are exactly identical
Sequences 2164.46 and 2164.50 are exactly identical
Sequences 2164.47 and 2164.48 are exactly identical
Sequences 39441.7 and 39441.8 are exactly identical

### Genome Statistics

| GenomeId | Total Genes | Single Copy | Used | Name |
|---|---|---|---|---|
| 190974.10 | 13 | 5 | 1 | Methanobrevibacter gottschalkii GG03_MAG_00003 |
| 1945577.3 | 738 | 248 | 80 | Methanosphaera sp. rholeuAM270 |
| 1945579.3 | 893 | 284 | 94 | Methanosphaera sp. rholeuAM74 |
| 2666342.12 | 901 | 298 | 96 | Methanosphaera sp. strain U.4_8 |
| 2666342.13 | 901 | 298 | 96 | Methanosphaera sp. U.4_8 |
| 2172.40 | 932 | 228 | 72 | Methanobrevibacter NG03_MAG_00264 |
| 2006182.3 | 953 | 305 | 97 | Methanobrevibacter sp. NOE |
| 1915625.3 | 953 | 269 | 86 | Methanobacteriaceae archaeon UBA588 |
| 1945578.3 | 961 | 305 | 98 | Methanosphaera sp. rholeuAM130 |
| 2099680.4 | 984 | 307 | 99 | Methanobacteriaceae archaeon strain SpSt-860 |
| 2099680.19 | 985 | 316 | 100 | Methanobacteriaceae archaeon strain MAG 290 |
| 2099680.21 | 985 | 316 | 100 | Methanobacteriaceae archaeon MAG 290 |
| 66852.12 | 995 | 281 | 93 | Methanobrevibacter sp. strain SIG15 |
| 2099680.18 | 997 | 284 | 94 | Methanobacteriaceae archaeon strain MAG 289 |
| 2099680.20 | 997 | 284 | 94 | Methanobacteriaceae archaeon MAG 289 |
| 1811676.3 | 1013 | 251 | 84 | Methanobacterium sp. PtaU1.Bin242 |
| 2317.19 | 1035 | 299 | 98 | Methanosphaera stadtmanae strain SIG23 |
| 1635284.4 | 1052 | 309 | 99 | Methanobacteriaceae archaeon 41_258 |
| 1945580.3 | 1071 | 314 | 98 | Methanosphaera sp. rholeuAM6 |
| 1789762.3 | 1076 | 317 | 100 | Methanosphaera sp. BMS |
| 2164.14 | 1078 | 268 | 84 | Methanobacterium sp. strain fen_5 |
| 2164.11 | 1084 | 255 | 87 | Methanobacterium sp. strain fen_9 |
| 2317.18 | 1100 | 314 | 99 | Methanosphaera stadtmanae strain SIG24 |
| 2164.7 | 1103 | 271 | 84 | Methanobacterium sp. strain fen_67 |
| 2317.20 | 1107 | 318 | 100 | Methanosphaera stadtmanae strain SIG25 |
| 1915624.3 | 1112 | 284 | 93 | Methanobacteriaceae archaeon UBA587 |
| 1561964.5 | 1144 | 318 | 99 | Methanosphaera sp. WGK6 |
| 66852.19 | 1152 | 282 | 93 | Methanobrevibacter sp. strain Rep1_Bin035 |
| 1915490.3 | 1158 | 224 | 72 | Methanobrevibacter sp. UBA594 |
| 2317.21 | 1161 | 295 | 93 | Methanosphaera stadtmanae strain ERR260183-bin.3 |
| 145262.18 | 1171 | 258 | 81 | Methanothermobacter thermautotrophicus strain UBA12524 |
| 2164.51 | 1199 | 307 | 95 | Methanobacterium sp. bin.336 |
| 190975.7 | 1214 | 293 | 94 | Methanobrevibacter thaueri strain SIG19 |
| 2164.48 | 1217 | 304 | 94 | Methanobacterium sp. MAG 286 |
| 2164.47 | 1217 | 304 | 94 | Methanobacterium sp. strain MAG 286 |
| 2172.49 | 1285 | 318 | 100 | Methanobrevibacter PG07_MAG_00003 |
| 66852.10 | 1296 | 309 | 97 | Methanobrevibacter sp. strain SIG16 |
| 230361.11 | 1296 | 285 | 91 | Methanobrevibacter millerae strain SIG13 |
| 66852.8 | 1316 | 288 | 89 | Methanobrevibacter sp. strain SIG22 |
| 2164.50 | 1317 | 300 | 97 | Methanobacterium sp. MAG 287 |
| 2164.46 | 1317 | 300 | 97 | Methanobacterium sp. strain MAG 287 |
| 1945576.3 | 1318 | 318 | 100 | Methanosphaera sp. DEW79 |
| 2164.12 | 1322 | 320 | 100 | Methanobacterium sp. strain fen_8 |
| 2164.13 | 1324 | 315 | 99 | Methanobacterium sp. strain fen_4 |
| 1860157.3 | 1325 | 306 | 98 | Methanosphaera sp. A6 |
| 190975.6 | 1352 | 306 | 97 | Methanobrevibacter thaueri strain SIG17 |
| 2317.11 | 1357 | 319 | 100 | Methanosphaera stadtmanae strain PA5 |

| | | | | |
|---|---|---|---|---|
| 2164.28 | 1378 | 260 | 86 | Methanobacterium sp. strain NC_groundwater_1554_Pr4_S-0.65um_39_6 |
| 294671.11 | 1390 | 311 | 100 | Methanobrevibacter olleyae strain SIG14 |
| 145262.19 | 1396 | 284 | 92 | Methanothermobacter thermautotrophicus strain UBA12174 |
| 1811674.3 | 1407 | 320 | 100 | Methanobacterium sp. PtaB.Bin024 |
| 66852.18 | 1411 | 312 | 100 | Methanobrevibacter sp. strain Rep2_Bin008 |
| 190975.5 | 1415 | 305 | 97 | Methanobrevibacter thaueri strain SIG18 |
| 230361.8 | 1422 | 320 | 100 | Methanobrevibacter millerae strain DSM 16643 |
| 2173.904 | 1437 | 318 | 100 | Methanobrevibacter smithii SUG1019 |
| 66852.9 | 1437 | 294 | 92 | Methanobrevibacter sp. strain SIG21 |
| 2164.27 | 1438 | 315 | 98 | Methanobacterium sp. strain SPRUCE.2016.2.metabat2.56 |
| 230361.12 | 1439 | 318 | 100 | Methanobrevibacter millerae strain SIG12 |
| 66852.11 | 1445 | 320 | 100 | Methanobrevibacter sp. strain SIG20 |
| 2173.813 | 1445 | 320 | 100 | Methanobrevibacter smithii strain MAG004 |
| 39441.8 | 1446 | 304 | 98 | Methanobrevibacter arboriphilus MAG 291 |
| 39441.7 | 1446 | 304 | 98 | Methanobrevibacter arboriphilus strain MAG 291 |
| 190976.3 | 1456 | 320 | 100 | Methanobrevibacter woesei strain DSM 11979 |
| 2164.15 | 1475 | 294 | 92 | Methanobacterium sp. strain bog_71 |
| 2173.826 | 1514 | 320 | 100 | Methanobrevibacter smithii strain L1_008_000M1_dasL1_008_000M1_concoct_61 |
| 1860156.3 | 1529 | 312 | 99 | Methanobrevibacter sp. A54 |
| 2173.812 | 1536 | 317 | 100 | Methanobrevibacter smithii strain COPD437 |
| 294671.4 | 1540 | 314 | 100 | Methanobrevibacter olleyae strain DSM 16632 |
| 2173.817 | 1550 | 320 | 100 | Methanobrevibacter smithii strain ERR260234-bin.14 |
| 420247.28 | 1575 | 320 | 100 | Methanobrevibacter smithii ATCC 35061 |
| 2173.795 | 1602 | 320 | 100 | Methanobrevibacter smithii strain MGYG-HGUT-02446 |
| 2173.70 | 1619 | 320 | 100 | Methanobrevibacter smithii strain WWM1085 |
| 1860099.3 | 1648 | 314 | 98 | Methanobrevibacter sp. A27 |
| 2164.6 | 1653 | 313 | 100 | Methanobacterium sp. strain palsa_68 |
| 187420.15 | 1665 | 319 | 100 | Methanothermobacter thermautotrophicus str. Delta H |
| 1811675.3 | 1724 | 306 | 96 | Methanobacterium sp. PtaU1.Bin097 |
| 1300164.4 | 1753 | 319 | 100 | Methanobrevibacter arboriphilus JCM 13429 = DSM 1125 strain DH1 |
| 2164.10 | 1759 | 300 | 95 | Methanobacterium sp. strain fen_48 |
| 2164.29 | 1784 | 319 | 100 | Methanobacterium sp. strain NC_groundwater_1739_Pr3_B-0.1um_39_112 |
| 2164.18 | 1785 | 307 | 98 | Methanobacterium sp. strain bog_16 |
| 2164.8 | 1792 | 304 | 98 | Methanobacterium sp. strain fen_50 |
| 1609968.3 | 1798 | 320 | 100 | Methanobrevibacter sp. YE315 |
| 190975.3 | 1857 | 320 | 100 | Methanobrevibacter thaueri strain DSM 11995 |
| 2164.31 | 1894 | 320 | 100 | Methanobacterium sp. strain Rep1_Bin027 |
| 2164.17 | 1910 | 319 | 100 | Methanobacterium sp. strain bog_22 |
| 2164.16 | 1950 | 316 | 99 | Methanobacterium sp. strain bog_69 |
| 2164.49 | 1989 | 319 | 100 | Methanobacterium sp. MAG 288 |
| 2164.45 | 1989 | 319 | 100 | Methanobacterium sp. strain MAG 288 |
| 59277.6 | 2003 | 319 | 100 | Methanobacterium subterraneum strain HO_2020 |
| 2173.814 | 2013 | 223 | 68 | Methanobrevibacter smithii strain MGYG-HGUT-04687 |
| 2173.815 | 2013 | 223 | 68 | Methanobrevibacter smithii strain ERR1297821-bin.15 |
| 2164.9 | 2018 | 314 | 99 | Methanobacterium sp. strain bog_57 |
| 1641383.3 | 2046 | 312 | 98 | Methanobacterium sp. 42_16 |
| 1220534.3 | 2072 | 309 | 95 | Methanobacterium sp. Maddingley MBC34 |
| 2164.4 | 2114 | 308 | 97 | Methanobacterium sp. strain bog_37 |
| 2164.5 | 2136 | 312 | 99 | Methanobacterium sp. strain bog_42 |
| 2164.3 | 2198 | 317 | 99 | Methanobacterium sp. strain bog_82 |
| 1204725.3 | 2203 | 320 | 100 | Methanobacterium formicicum DSM 3637 |
| 59277.3 | 2227 | 319 | 100 | Methanobacterium subterraneum strain A8p |
| 1860100.3 | 2857 | 319 | 100 | Methanobacterium sp. A39 |

### Gene Family Statistics

Gene families are ranked by alignment score combining mean per-position variability, alignment length, and gappiness.

| PGFam | Align. Score | Align. Length | Num Seqs | Mean Sqr Freq | Prop Gaps | Used In Analysis | Product |
|---|---|---|---|---|---|---|---|
| PGF_01175575 | 15.09 | 630 | 99 | 0.601 | 0.069 | True | Threonyl-tRNA synthetase (EC 6.1.1.3) |
| PGF_05070366 | 14.92 | 573 | 96 | 0.623 | 0.077 | True | Dihydroxy-acid dehydratase (EC 4.2.1.9) |
| PGF_00018085 | 14.73 | 555 | 95 | 0.625 | 0.084 | True | Lysyl-tRNA synthetase (class I) (EC 6.1.1.6) |
| PGF_00065138 | 14.39 | 420 | 97 | 0.702 | 0.076 | True | Uncharacterized protein MJ0800 |
| PGF_00421761 | 14.24 | 321 | 96 | 0.795 | 0.054 | True | DNA repair and recombination protein RadA |
| PGF_00052238 | 14.05 | 454 | 99 | 0.659 | 0.070 | True | Signal recognition particle protein Ffh |
| PGF_03609651 | 14.00 | 720 | 97 | 0.522 | 0.148 | True | Methionyl-tRNA synthetase (EC 6.1.1.10) |
| PGF_00424924 | 13.73 | 418 | 94 | 0.672 | 0.047 | True | Eukaryotic peptide chain release factor subunit 1 |
| PGF_00012928 | 13.36 | 378 | 92 | 0.687 | 0.106 | True | Hydroxymethylglutaryl-CoA synthase (EC 2.3.3.10) |
| PGF_00426105 | 13.26 | 439 | 97 | 0.633 | 0.129 | True | 5,6,7,8-tetrahydromethanopterin hydro-lyase (EC 4.2.1.147) @ Formaldehyde activating enzyme / D-arabino-3-hexulose 6-phosphate formaldehyde-lyase (EC 4.1.2.43) |
| PGF_00012505 | 13.25 | 492 | 97 | 0.597 | 0.097 | True | Hydrogenase expression/formation protein |
| PGF_10373476 | 13.25 | 442 | 97 | 0.630 | 0.067 | True | Diaminopimelate decarboxylase (EC 4.1.1.20) |
| PGF_09019918 | 13.24 | 436 | 92 | 0.634 | 0.043 | True | Glutamate-1-semialdehyde 2,1-aminomutase (EC 5.4.3.8) |
| PGF_03316046 | 13.23 | 464 | 96 | 0.614 | 0.176 | True | Threonine synthase (EC 4.2.3.1) |
| PGF_00012926 | 13.23 | 413 | 93 | 0.651 | 0.042 | True | Hydroxymethylglutaryl-CoA reductase (EC 1.1.1.34) |
| PGF_00015813 | 13.22 | 722 | 98 | 0.492 | 0.176 | True | Uncharacterized KH and PIN-domain containing protein MJ1533 |
| PGF_03590193 | 13.18 | 445 | 98 | 0.625 | 0.087 | True | Serine hydroxymethyltransferase (tetrahydromethanopterin-dependent) |
| PGF_02620298 | 13.10 | 484 | 93 | 0.595 | 0.058 | True | Argininosuccinate lyase (EC 4.3.2.1) |
| PGF_07750515 | 13.08 | 455 | 95 | 0.613 | 0.067 | True | Phosphoribosylamine--glycine ligase (EC 6.3.4.13) |
| PGF_00065212 | 12.74 | 429 | 96 | 0.615 | 0.076 | True | Uncharacterized protein MJ1313 |
| PGF_00006244 | 12.73 | 387 | 98 | 0.647 | 0.107 | True | Formate--phosphoribosylaminoimidazolecarboxamide ligase (EC 6.3.4.23) |
| PGF_00064425 | 12.57 | 453 | 95 | 0.591 | 0.098 | True | UbiD family decarboxylase, MJ1133 type |
| PGF_00008338 | 12.51 | 463 | 95 | 0.581 | 0.082 | True | Glutamyl-tRNA(Gln) amidotransferase asparaginase subunit (EC 6.3.5.7) |
| PGF_03060918 | 12.40 | 406 | 93 | 0.615 | 0.064 | True | Uncharacterized protein SSO2743 |
| PGF_04375438 | 12.28 | 420 | 94 | 0.599 | 0.115 | True | Coenzyme B synthesis from 2-oxoglutarate: steps 1, 6, and 10 |
| PGF_00421409 | 12.18 | 724 | 94 | 0.453 | 0.156 | True | DNA mismatch repair protein |
| PGF_00046029 | 12.09 | 293 | 91 | 0.707 | 0.059 | True | Pyruvate:ferredoxin oxidoreductase, beta subunit (EC 1.2.7.1) |
| PGF_00067188 | 11.80 | 363 | 96 | 0.619 | 0.092 | True | Aspartate-semialdehyde dehydrogenase (EC 1.2.1.11) |
| PGF_00024323 | 11.76 | 364 | 98 | 0.617 | 0.119 | True | NAD(P)-dependent glyceraldehyde-3-phosphate dehydrogenase, archaeal (EC 1.2.1.59) |
| PGF_00006981 | 11.76 | 335 | 95 | 0.643 | 0.072 | True | GTP cyclohydrolase MptA (EC 3.5.4.39) |
| PGF_10350547 | 11.71 | 396 | 95 | 0.589 | 0.092 | True | 3-dehydroquinate synthase II (EC 1.4.1.24) |
| PGF_00121834 | 11.59 | 420 | 93 | 0.565 | 0.055 | True | hypothetical protein |
| PGF_08454293 | 11.58 | 459 | 97 | 0.541 | 0.113 | True | N-acetylglucosamine-1-phosphate uridyltransferase (EC 2.7.7.23) / Glucosamine-1-phosphate N-acetyltransferase (EC 2.3.1.157) |
| PGF_00067772 | 11.58 | 405 | 98 | 0.575 | 0.185 | True | [NiFe] hydrogenase metallocenter assembly protein HypE |
| PGF_02474526 | 11.56 | 348 | 95 | 0.619 | 0.061 | True | anion-transporting ATPase |
| PGF_03375688 | 11.55 | 340 | 95 | 0.626 | 0.109 | True | S-methyl-5-thioribose-1-phosphate isomerase (EC 5.3.1.23) |
| PGF_00424971 | 11.48 | 783 | 95 | 0.410 | 0.269 | True | Excinuclease ABC subunit C |
| PGF_02277678 | 11.43 | 423 | 95 | 0.556 | 0.093 | True | Phosphoglycerate kinase (EC 2.7.2.3) |
| PGF_00033954 | 11.33 | 366 | 95 | 0.592 | 0.095 | True | Phosphoribosylformylglycinamidine cyclo-ligase (EC 6.3.3.1) |
| PGF_02961521 | 11.31 | 656 | 96 | 0.442 | 0.252 | True | Fumarate reductase (CoM/CoB), subunit TfrB (EC 1.3.4.1) |
| PGF_00015737 | 11.31 | 372 | 95 | 0.586 | 0.093 | True | Isopentenyl-diphosphate delta-isomerase, FMN-dependent (EC 5.3.3.2) |
| PGF_00413256 | 11.29 | 454 | 91 | 0.530 | 0.088 | True | tRNA N6-threonylcarbamoyladenosine 2-methylthiotransferase (putative) @ tRNA-i(6)A37 methylthiotransferase (EC 2.8.4.3) |
| PGF_00853393 | 11.23 | 302 | 95 | 0.646 | 0.078 | True | Succinyl-CoA ligase [ADP-forming] alpha chain (EC 6.2.1.5) |
| PGF_00049910 | 11.18 | 202 | 91 | 0.786 | 0.028 | True | SSU ribosomal protein SAe (S2p) |
| PGF_00413286 | 11.16 | 540 | 96 | 0.480 | 0.238 | True | tRNA pseudouridine(13) synthase (EC 5.4.99.27) |
| PGF_00424925 | 11.13 | 274 | 93 | 0.672 | 0.075 | True | Eukaryotic translation initiation factor 2 alpha subunit |
| PGF_00006318 | 11.05 | 319 | 96 | 0.619 | 0.094 | True | Formylmethanofuran--tetrahydromethanopterin N-formyltransferase (EC 2.3.1.101) |
| PGF_06674514 | 11.00 | 309 | 94 | 0.626 | 0.090 | True | Nitrogenase FeMo-cofactor synthesis FeS core scaffold and assembly protein NifB |

| | | | | | | | |
|---|---|---|---|---|---|---|---|
| PGF_00070777 | 10.89 | 344 | 97 | 0.587 | 0.043 | True | Diphthamide biosynthesis protein 1 |
| PGF_03004613 | 10.87 | 495 | 96 | 0.488 | 0.161 | True | Histidyl-tRNA synthetase (EC 6.1.1.21) |
| PGF_00005666 | 10.83 | 345 | 93 | 0.583 | 0.057 | True | Flap structure-specific endonuclease |
| PGF_01087653 | 10.82 | 314 | 95 | 0.611 | 0.179 | True | 2-amino-3,7-dideoxy-D-threo-hept-6-ulosonate synthase (EC 2.2.1.10) |
| PGF_07015581 | 10.73 | 426 | 96 | 0.520 | 0.184 | True | Carbamoyl-phosphate synthase small chain (EC 6.3.5.5) |
| PGF_00489714 | 10.69 | 369 | 95 | 0.557 | 0.139 | True | Porphobilinogen synthase (EC 4.2.1.24) |
| PGF_00063915 | 10.69 | 336 | 96 | 0.583 | 0.090 | True | Tyrosyl-tRNA synthetase (EC 6.1.1.1) |
| PGF_00026805 | 10.67 | 334 | 96 | 0.584 | 0.102 | True | OB-fold nucleic acid binding domain protein |
| PGF_06230969 | 10.63 | 565 | 94 | 0.447 | 0.130 | True | NAD(P)H-hydrate epimerase (EC 5.1.99.6) / ADP-dependent (S)-NAD(P)H-hydrate dehydratase (EC 4.2.1.136) |
| PGF_00063594 | 10.63 | 673 | 96 | 0.410 | 0.252 | True | Archaeal DNA polymerase II small subunit (EC 2.7.7.7) |
| PGF_08376928 | 10.62 | 344 | 95 | 0.573 | 0.133 | True | Branched-chain amino acid aminotransferase (EC 2.6.1.42) |
| PGF_02011315 | 10.62 | 349 | 94 | 0.568 | 0.077 | True | Geranylgeranyl diphosphate synthase (EC 2.5.1.29) |
| PGF_00421629 | 10.56 | 249 | 96 | 0.669 | 0.049 | True | DNA polymerase sliding clamp protein PCNA |
| PGF_00413307 | 10.56 | 524 | 93 | 0.461 | 0.209 | True | tRNA(Ile)(2)-agmatinylcytidine synthase (EC 6.3.4.22) |
| PGF_00549380 | 10.49 | 300 | 95 | 0.606 | 0.034 | True | 4-hydroxy-tetrahydrodipicolinate synthase (EC 4.3.3.7) |
| PGF_00424417 | 10.46 | 359 | 92 | 0.552 | 0.094 | True | Energy conserving hydrogenase Ehb integral membrane protein O |
| PGF_00008548 | 10.45 | 438 | 96 | 0.499 | 0.219 | True | Glycerol-1-phosphate dehydrogenase [NAD(P)+] (EC 1.1.1.261) |
| PGF_12766951 | 10.38 | 386 | 94 | 0.529 | 0.089 | True | Phospho-N-acetylmuramoyl-pentapeptide-transferase (EC 2.7.8.13) |
| PGF_12884281 | 10.37 | 428 | 98 | 0.501 | 0.117 | True | dNTP triphosphohydrolase, broad substrate specificity |
| PGF_00413206 | 10.33 | 452 | 94 | 0.486 | 0.155 | True | tRNA (guanine(26)-N(2))-dimethyltransferase (EC 2.1.1.216) |
| PGF_00008334 | 10.31 | 433 | 92 | 0.495 | 0.115 | True | Glutamyl-tRNA reductase (EC 1.2.1.70) |
| PGF_00000125 | 10.30 | 384 | 97 | 0.526 | 0.079 | True | 7,8-didemethyl-8-hydroxy-5-deazariboflavin synthase subunit 1 |
| PGF_00421665 | 10.27 | 581 | 95 | 0.426 | 0.340 | True | DNA primase, archaeal DnaG-type |
| PGF_07889681 | 10.21 | 387 | 95 | 0.519 | 0.164 | True | N-acetyl-gamma-glutamyl-phosphate reductase (EC 1.2.1.38) |
| PGF_00424387 | 10.20 | 320 | 93 | 0.570 | 0.091 | True | Energy conserving hydrogenase Eha associated protein (protein S); Formylmethanofuran--tetrahydromethanopterin N-formyltransferase (EC 2.3.1.101) |
| PGF_00048585 | 10.16 | 262 | 94 | 0.627 | 0.113 | True | Exosome complex exonuclease RRP41 |
| PGF_00016613 | 10.12 | 328 | 91 | 0.559 | 0.093 | True | Lactyl (2) diphospho-(5')guanosine:7,8-didemethyl-8-hydroxy-5-deazariboflavin 2-phospho-L-lactate transferase (EC 2.7.8.28) |
| PGF_02960239 | 10.10 | 459 | 93 | 0.471 | 0.071 | True | TldE protein, part of TldE/TldD proteolytic complex |
| PGF_00033959 | 10.09 | 226 | 93 | 0.671 | 0.058 | True | Phosphoribosylformylglycinamidine synthase, glutamine amidotransferase subunit (EC 6.3.5.3) |
| PGF_00064367 | 10.01 | 409 | 95 | 0.495 | 0.093 | True | UPF0425 pyridoxal phosphate-dependent protein MJ0158 |
| PGF_00051879 | 9.87 | 234 | 95 | 0.645 | 0.034 | True | Shwachman-Bodian-Diamond syndrome protein; predicted exosome subunit |
| PGF_02329487 | 9.87 | 364 | 90 | 0.517 | 0.074 | True | 4-[[4-(2-aminoethyl)phenoxy]-methyl]-2-furanmethanamine-glutamate synthase |
| PGF_10097367 | 9.81 | 319 | 93 | 0.549 | 0.105 | True | Porphobilinogen deaminase (EC 2.5.1.61) |
| PGF_00418425 | 9.66 | 365 | 93 | 0.506 | 0.215 | True | Coenzyme F420 hydrogenase beta subunit (FrcB) (EC 1.12.98.1) |
| PGF_00415574 | 9.62 | 271 | 93 | 0.584 | 0.061 | True | FIG00137952: Uncharacterized protein (ATP-grasp superfamily) |
| PGF_00065257 | 9.54 | 273 | 91 | 0.577 | 0.118 | True | Uncharacterized protein MJ1676 |
| PGF_00016267 | 9.48 | 573 | 97 | 0.396 | 0.339 | True | L-tyrosine decarboxylase (EC 4.1.1.25) |
| PGF_02281324 | 9.41 | 424 | 92 | 0.457 | 0.188 | True | Uncharacterized protein MJ1086 |
| PGF_00034252 | 9.39 | 446 | 95 | 0.445 | 0.142 | True | 2,5-diamino-6-ribitylamino-pyrimidinone 5-phosphate deaminase, archaeal (EC 3.5.4.-) |
| PGF_02346669 | 9.37 | 358 | 97 | 0.495 | 0.179 | True | Ornithine carbamoyltransferase (EC 2.1.3.3) |
| PGF_00424431 | 9.28 | 166 | 90 | 0.720 | 0.114 | True | Energy conserving hydrogenase Ehb small subunit (protein M) |
| PGF_00853991 | 9.23 | 312 | 97 | 0.522 | 0.106 | True | Diaminopimelate epimerase (EC 5.1.1.7) |
| PGF_00414927 | 9.18 | 560 | 93 | 0.388 | 0.197 | True | CCA tRNA nucleotidyltransferase, archaeal type (EC 2.7.7.72) |
| PGF_00048782 | 9.16 | 361 | 93 | 0.482 | 0.202 | True | Ribose-phosphate pyrophosphokinase (EC 2.7.6.1) |
| PGF_00065013 | 9.16 | 485 | 94 | 0.416 | 0.393 | True | Uncharacterized protein MJ0094 |
| PGF_00424430 | 9.12 | 239 | 91 | 0.590 | 0.094 | True | Energy conserving hydrogenase Ehb protein Q |
| PGF_00037061 | 9.07 | 275 | 95 | 0.547 | 0.069 | True | Exosome complex exonuclease RRP45 |
| PGF_00418427 | 9.00 | 335 | 94 | 0.492 | 0.233 | True | Coenzyme F420 hydrogenase gamma subunit (FrcG) (EC 1.12.98.1) |
| PGF_01513745 | 8.90 | 243 | 92 | 0.571 | 0.154 | True | Proteasome subunit beta (EC 3.4.25.1), archaeal |
| PGF_00422341 | 8.90 | 211 | 94 | 0.612 | 0.118 | True | DNA-directed RNA polymerase subunit E' (EC 2.7.7.6) |
| PGF_00260942 | 8.89 | 474 | 90 | 0.408 | 0.252 | True | UPF0348 protein MJ0951 |
| PGF_05657253 | 8.84 | 245 | 96 | 0.565 | 0.099 | True | Ribose-5-phosphate isomerase A (EC 5.3.1.6) |

| | | | | | | | |
|---|---|---|---|---|---|---|---|
| PGF_00016341 | 8.71 | 215 | 95 | 0.594 | 0.184 | False | LSU ribosomal protein L15e |
| PGF_00421762 | 8.71 | 239 | 92 | 0.564 | 0.032 | False | DNA repair and recombination protein RadB |
| PGF_00013803 | 8.71 | 199 | 95 | 0.618 | 0.037 | False | Imidazoleglycerol-phosphate dehydratase (EC 4.2.1.19) |
| PGF_00049839 | 8.59 | 133 | 91 | 0.745 | 0.013 | False | SSU ribosomal protein S13e (S15p) |
| PGF_00411462 | 8.53 | 235 | 94 | 0.557 | 0.088 | False | Box C/D RNA-guided RNA methyltransferase subunit fibrillarin |
| PGF_00413313 | 8.49 | 289 | 92 | 0.500 | 0.160 | False | tRNA(Phe) (4-demethylwyosine(37)-C(7)) aminocarboxypropyltransferase (EC 2.5.1.114) |
| PGF_00625415 | 8.48 | 255 | 94 | 0.531 | 0.123 | False | 7-cyano-7-deazaguanine synthase (EC 6.3.4.20) |
| PGF_07097873 | 8.48 | 351 | 94 | 0.453 | 0.186 | False | Ribonuclease Z (EC 3.1.26.11) |
| PGF_01219411 | 8.47 | 350 | 91 | 0.453 | 0.094 | False | 5-methyltetrahydropteroyltriglutamate--homocysteine methyltransferase (EC 2.1.1.14) |
| PGF_00413205 | 8.45 | 417 | 97 | 0.414 | 0.191 | False | tRNA (guanine(10)-N(2))-dimethyltransferase (EC 2.1.1.213) |
| PGF_00066972 | 8.41 | 203 | 93 | 0.590 | 0.112 | False | Xanthosine/inosine triphosphate pyrophosphatase |
| PGF_00413210 | 8.41 | 436 | 95 | 0.403 | 0.239 | False | tRNA (guanine(37)-N(1))-methyltransferase (EC 2.1.1.228) |
| PGF_00415581 | 8.39 | 224 | 97 | 0.561 | 0.085 | False | 23S rRNA (uridine(2552)-2'-O)-methyltransferase (EC 2.1.1.166) |
| PGF_08473931 | 8.39 | 199 | 96 | 0.595 | 0.077 | False | Flavin prenyltransferase UbiX |
| PGF_02283727 | 8.37 | 145 | 98 | 0.695 | 0.085 | False | Eukaryotic translation initiation factor 2 beta subunit |
| PGF_00049827 | 8.32 | 335 | 98 | 0.454 | 0.158 | False | SSU rRNA (adenine(1518)-N(6)/adenine(1519)-N(6))-dimethyltransferase (EC 2.1.1.182) |
| PGF_00418292 | 8.31 | 528 | 97 | 0.362 | 0.310 | False | Cobalt-precorrin-5B (C1)-methyltransferase (EC 2.1.1.195) |
| PGF_02925426 | 8.30 | 541 | 92 | 0.357 | 0.276 | False | Uncharacterized protein MJ0971 |
| PGF_00426932 | 8.27 | 149 | 94 | 0.677 | 0.108 | False | 6,7-dimethyl-8-ribityllumazine synthase (EC 2.5.1.78) |
| PGF_00048479 | 8.25 | 163 | 92 | 0.647 | 0.097 | False | Riboflavin synthase archaeal (EC 2.5.1.9) |
| PGF_08181546 | 8.24 | 346 | 91 | 0.443 | 0.201 | False | Shikimate 5-dehydrogenase I alpha (EC 1.1.1.25) |
| PGF_00051543 | 8.21 | 381 | 96 | 0.421 | 0.245 | False | Shikimate kinase II (EC 2.7.1.71) |
| PGF_00411463 | 8.15 | 462 | 92 | 0.379 | 0.196 | False | Box C/D RNA-guided RNA methyltransferase subunit Nop5 |
| PGF_00025580 | 8.04 | 190 | 97 | 0.583 | 0.095 | False | Nicotinamide-nucleotide adenylyltransferase, NadM family (EC 2.7.7.1) |
| PGF_00070158 | 8.04 | 133 | 96 | 0.697 | 0.025 | False | conserved protein associated with acetyl-CoA C-acyltransferase and HMGCo |
| PGF_01435183 | 8.02 | 203 | 96 | 0.563 | 0.078 | False | Adenine phosphoribosyltransferase (EC 2.4.2.7) |
| PGF_03518570 | 8.01 | 170 | 95 | 0.614 | 0.136 | False | Nucleoside diphosphate kinase (EC 2.7.4.6) |
| PGF_07544077 | 8.00 | 324 | 95 | 0.444 | 0.192 | False | Prephenate dehydratase (EC 4.2.1.51) |
| PGF_04867467 | 7.99 | 222 | 94 | 0.536 | 0.103 | False | Imidazole glycerol phosphate synthase amidotransferase subunit HisH |
| PGF_00013510 | 7.94 | 236 | 93 | 0.517 | 0.145 | False | IMP cyclohydrolase (EC 3.5.4.10) [alternate form] |
| PGF_00015730 | 7.91 | 334 | 93 | 0.433 | 0.222 | False | Isopentenyl phosphate kinase (EC 2.7.4.26) |
| PGF_00049978 | 7.74 | 253 | 93 | 0.486 | 0.117 | False | 2-amino-5-formylamino-6-ribosylaminopyrimidin-4(3H)-one 5-monophosphate deformylase (EC 3.5.1.102) |
| PGF_00006991 | 7.57 | 532 | 95 | 0.328 | 0.529 | False | GTP cyclohydrolase III (EC 3.5.4.29) |
| PGF_00061926 | 7.42 | 364 | 94 | 0.389 | 0.225 | False | Triphosphoribosyl-dephospho-CoA synthetase |
| PGF_00416211 | 7.40 | 367 | 93 | 0.386 | 0.256 | False | Candidate phosphomevalonate decarboxylase; COG1355, Predicted dioxygenase |
| PGF_00065006 | 7.37 | 744 | 97 | 0.270 | 0.490 | False | Uncharacterized protein MJ0065 |
| PGF_00066960 | 7.32 | 181 | 91 | 0.544 | 0.139 | False | Aspartate carbamoyltransferase regulatory chain (PyrI) |
| PGF_00016426 | 7.30 | 91 | 92 | 0.765 | 0.029 | False | LSU ribosomal protein L37Ae |
| PGF_00418309 | 7.27 | 280 | 95 | 0.434 | 0.279 | False | Cobalt-precorrin-8 methylmutase (EC 5.4.99.60) |
| PGF_00060434 | 7.21 | 127 | 93 | 0.640 | 0.188 | False | Translation initiation factor 1A |
| PGF_02781328 | 7.21 | 338 | 91 | 0.392 | 0.232 | False | NAD synthetase (EC 6.3.1.5) |
| PGF_02278091 | 7.16 | 423 | 94 | 0.348 | 0.401 | False | Serine/threonine-protein kinase RIO1 (EC 2.7.11.1) |
| PGF_00016438 | 7.11 | 112 | 91 | 0.672 | 0.183 | False | LSU ribosomal protein L44e |
| PGF_00070870 | 7.08 | 244 | 95 | 0.453 | 0.199 | False | eRF1 methyltransferase catalytic subunit MTQ2 |
| PGF_02278032 | 7.05 | 219 | 91 | 0.476 | 0.120 | False | Uncharacterized C4-type Zn finger protein MJ0530 |
| PGF_00424939 | 7.04 | 169 | 92 | 0.542 | 0.252 | False | Eukaryotic translation initiation factor 5A |
| PGF_02191019 | 7.02 | 486 | 95 | 0.318 | 0.320 | False | Thiamine-monophosphate kinase (EC 2.7.4.16) |
| PGF_00413290 | 6.96 | 409 | 91 | 0.344 | 0.325 | False | tRNA pseudouridine(38-40) synthase (EC 5.4.99.12) |
| PGF_00055077 | 6.92 | 230 | 90 | 0.456 | 0.206 | False | Sulfopyruvate decarboxylase - beta subunit (EC 4.1.1.79) |
| PGF_00413204 | 6.85 | 249 | 98 | 0.434 | 0.288 | False | tRNA (cytidine(56)-2'-O)-methyltransferase (EC 2.1.1.206) |
| PGF_00049898 | 6.80 | 149 | 94 | 0.557 | 0.158 | False | SSU ribosomal protein S6e |
| PGF_00016363 | 6.77 | 99 | 96 | 0.681 | 0.031 | False | LSU ribosomal protein L21e |
| PGF_00056896 | 6.66 | 271 | 91 | 0.405 | 0.196 | False | Thymidylate synthase (EC 2.1.1.45) |
| PGF_00049905 | 6.65 | 141 | 90 | 0.560 | 0.134 | False | SSU ribosomal protein S8e |
| [illegible] | [illegible] | [illegible] | [illegible] | [illegible] | [illegible] | [illegible] | [illegible] |

| | | | | | | | |
|---|---|---|---|---|---|---|---|
| PGF_04359948 | 6.61 | 864 | 96 | 0.225 | 0.620 | False | Selenophosphate synthetase-related protein MJ0640 |
| PGF_00424416 | 6.54 | 229 | 92 | 0.432 | 0.279 | False | Energy conserving hydrogenase Ehb ferredoxin-containing protein L |
| PGF_01513918 | 6.51 | 224 | 93 | 0.435 | 0.193 | False | Adenylate cyclase (EC 4.6.1.1) |
| PGF_04815798 | 6.39 | 630 | 96 | 0.255 | 0.492 | False | Mevalonate kinase (EC 2.7.1.36) |
| PGF_00049859 | 6.35 | 242 | 90 | 0.408 | 0.407 | False | SSU ribosomal protein S19e |
| PGF_02776739 | 6.29 | 161 | 95 | 0.496 | 0.181 | False | Phosphoribosyl-AMP cyclohydrolase (EC 3.5.4.19) |
| PGF_00065054 | 6.14 | 141 | 92 | 0.517 | 0.139 | False | Uncharacterized protein MJ0408 |
| PGF_00046039 | 6.09 | 201 | 92 | 0.430 | 0.268 | False | Pyruvoyl-dependent arginine decarboxylase (EC 4.1.1.19) |
| PGF_00065076 | 6.07 | 219 | 94 | 0.410 | 0.301 | False | Uncharacterized protein MJ0498 |
| PGF_00083887 | 5.84 | 129 | 91 | 0.515 | 0.307 | False | hypothetical protein |
| PGF_00016417 | 5.83 | 115 | 90 | 0.544 | 0.234 | False | LSU ribosomal protein L34e |
| PGF_00067769 | 5.80 | 92 | 95 | 0.604 | 0.099 | False | [NiFe] hydrogenase metallocenter assembly protein HypC |
| PGF_02976460 | 5.78 | 380 | 91 | 0.297 | 0.494 | False | Phosphatidylinositol phosphate synthase @ Archaetidylinositol phosphate synthase (EC 2.7.8.39) |
| PGF_00067788 | 5.73 | 670 | 92 | 0.222 | 0.674 | False | [NiFe] hydrogenase nickel incorporation-associated protein HypB |
| PGF_07957805 | 5.60 | 93 | 90 | 0.581 | 0.086 | False | Phosphoribosylformylglycinamidine synthase, PurS subunit (EC 6.3.5.3) |
| PGF_02278971 | 3.47 | 296 | 93 | 0.202 | 0.668 | False | PaaD-like protein MJ1129 |

### Phylogenetic Tree Report

#### Rendered Tree

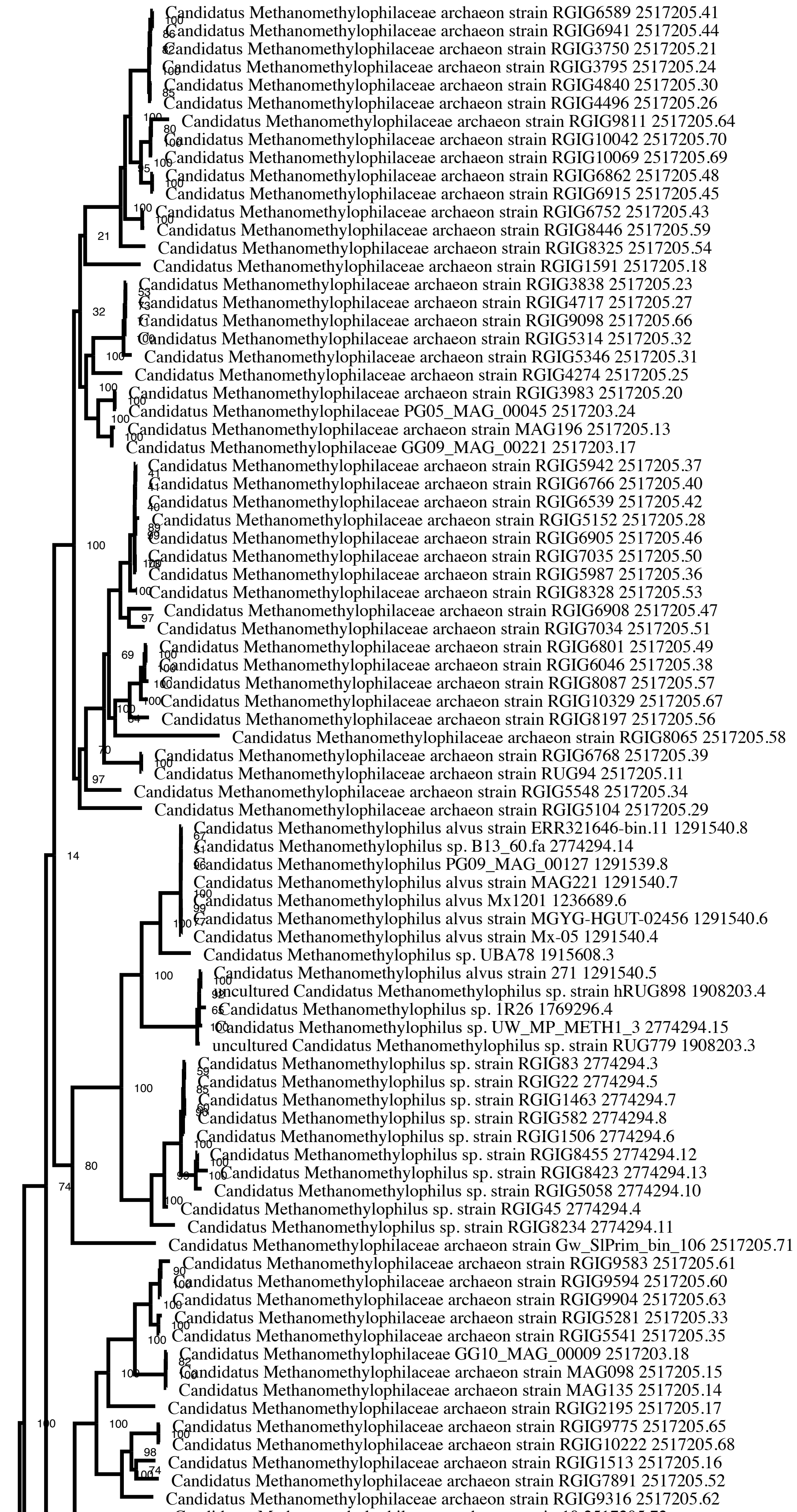

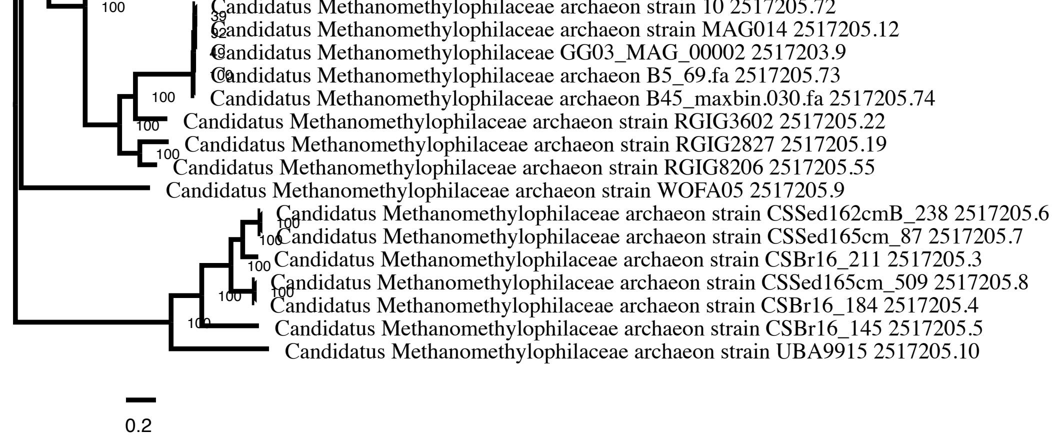


[Alternate View]

[Files with more details]

### Tree Analysis Statistics

| | |
|---|---|
| **Requested genomes** | 100 |
| **Genomes with data** | 99 |
| **Genomes lacking data** | 1 |
| **Max allowed deletions** | 10 |
| **Max allowed duplications** | 10 |
| **Single-copy genes requested** | 100 |
| **Single-copy genes found** | 7 |
| **Num protein alignments** | 7 |
| **Alignment program** | mafft |
| **Protein alignment time** | 145.2 seconds |
| **Num aligned amino acids** | 3041 |
| **Num CDS alignments** | 7 |
| **Num aligned nucleotides** | 9123 |
| **Best protein model found by RAxML** | LG |
| **Branch support method** | RAxML Fast Bootstrapping |
| **RAxML likelihood** | -258394.3039 |
| **RAxML version** | 8.2.11 |
| **RAxML time** | 9361.9 seconds |
| **Total time** | 9601.1 seconds |

### RAxML Command Line

Goal: Analyze proteins with model 'AUTO' to find best substitution model.
raxmlHPC-PTHREADS-SSE3 -s Methanomethylophilaceae_10_MAX_DELs_DUPs_proteins.phy -n Methanomethylophilaceae_10_MAX_DELs_DUPs_proteins -m PROTCATAUTO -p 12345 -T 12 -e 10
Process time: 1060.386 seconds

Goal: Find best tree.
raxmlHPC-PTHREADS-SSE3 -s Methanomethylophilaceae_10_MAX_DELs_DUPs.phy -n Methanomethylophilaceae_10_MAX_DELs_DUPs -m GTRCAT -q Methanomethylophilaceae_10_MAX_DELs_DUPs.partitions -p 12345 -T 12 -f a -x 12345 -N 100
Process time: 8301.561 seconds

### RAxML Codon and Amino Acid Partitions

```
DNA, codon1 = 1-9123\3
DNA, codon2 = 2-9123\3
DNA, codon3 = 3-9123\3
LG, proteins = 9124-12164
```

### Genome Statistics

| GenomeId | Total Genes | Single Copy | Used | Name |
|---|---|---|---|---|
| 2517205.71 | 177 | 4 | 4 | Candidatus Methanomethylophilaceae archaeon strain Gw_SlPrim_bin_106 |
| 2517205.60 | 193 | 5 | 5 | Candidatus Methanomethylophilaceae archaeon strain RGIG9594 |
| 2517205.19 | 260 | 7 | 7 | Candidatus Methanomethylophilaceae archaeon strain RGIG2827 |
| 2517205.59 | 261 | 5 | 5 | Candidatus Methanomethylophilaceae archaeon strain RGIG8446 |
| 2517205.52 | 263 | 7 | 7 | Candidatus Methanomethylophilaceae archaeon strain RGIG7891 |
| 2517205.67 | 266 | 4 | 4 | Candidatus Methanomethylophilaceae archaeon strain RGIG10329 |
| 2774294.8 | 270 | 6 | 6 | Candidatus Methanomethylophilus sp. strain RGIG582 |
| 2774294.10 | 270 | 7 | 7 | Candidatus Methanomethylophilus sp. strain RGIG5058 |
| 2517205.57 | 274 | 5 | 5 | Candidatus Methanomethylophilaceae archaeon strain RGIG8087 |
| 2517205.8 | 275 | 4 | 4 | Candidatus Methanomethylophilaceae archaeon strain CSSed165cm_509 |
| 2517205.33 | 277 | 5 | 5 | Candidatus Methanomethylophilaceae archaeon strain RGIG5281 |
| 2517205.65 | 288 | 5 | 5 | Candidatus Methanomethylophilaceae archaeon strain RGIG9775 |
| 2517205.68 | 291 | 5 | 5 | Candidatus Methanomethylophilaceae archaeon strain RGIG10222 |
| 2517205.64 | 293 | 6 | 6 | Candidatus Methanomethylophilaceae archaeon strain RGIG9811 |
| 2517205.63 | 294 | 7 | 7 | Candidatus Methanomethylophilaceae archaeon strain RGIG9904 |
| 2517205.55 | 307 | 6 | 6 | Candidatus Methanomethylophilaceae archaeon strain RGIG8206 |
| 2517205.30 | 311 | 5 | 5 | Candidatus Methanomethylophilaceae archaeon strain RGIG4840 |
| 2517205.31 | 314 | 6 | 6 | Candidatus Methanomethylophilaceae archaeon strain RGIG5346 |
| 2517205.28 | 317 | 6 | 6 | Candidatus Methanomethylophilaceae archaeon strain RGIG5152 |
| 2517205.62 | 323 | 7 | 7 | Candidatus Methanomethylophilaceae archaeon strain RGIG9316 |
| 2517205.61 | 326 | 6 | 6 | Candidatus Methanomethylophilaceae archaeon strain RGIG9583 |
| 2774294.3 | 329 | 6 | 6 | Candidatus Methanomethylophilus sp. strain RGIG83 |
| 2517205.7 | 330 | 4 | 4 | Candidatus Methanomethylophilaceae archaeon strain CSSed165cm_87 |
| 2517205.3 | 331 | 7 | 7 | Candidatus Methanomethylophilaceae archaeon strain CSBr16_211 |
| 2517205.56 | 334 | 6 | 6 | Candidatus Methanomethylophilaceae archaeon strain RGIG8197 |
| 2517205.21 | 334 | 5 | 5 | Candidatus Methanomethylophilaceae archaeon strain RGIG3750 |
| 2517205.20 | 335 | 5 | 5 | Candidatus Methanomethylophilaceae archaeon strain RGIG3983 |
| 2517205.50 | 355 | 7 | 7 | Candidatus Methanomethylophilaceae archaeon strain RGIG7035 |
| 2517205.5 | 356 | 3 | 3 | Candidatus Methanomethylophilaceae archaeon strain CSBr16_145 |
| 2517205.10 | 361 | 7 | 7 | Candidatus Methanomethylophilaceae archaeon strain UBA9915 |
| 2517205.9 | 364 | 5 | 5 | Candidatus Methanomethylophilaceae archaeon strain WOFA05 |
| 2517205.69 | 366 | 6 | 6 | Candidatus Methanomethylophilaceae archaeon strain RGIG10069 |
| 2517205.23 | 367 | 6 | 6 | Candidatus Methanomethylophilaceae archaeon strain RGIG3838 |
| 2517203.17 | 374 | 6 | 6 | Candidatus Methanomethylophilaceae GG09_MAG_00221 |
| 2517205.40 | 377 | 5 | 5 | Candidatus Methanomethylophilaceae archaeon strain RGIG6766 |
| 2517205.53 | 380 | 6 | 6 | Candidatus Methanomethylophilaceae archaeon strain RGIG8328 |
| 2517205.70 | 383 | 6 | 6 | Candidatus Methanomethylophilaceae archaeon strain RGIG10042 |
| 2517205.49 | 397 | 7 | 7 | Candidatus Methanomethylophilaceae archaeon strain RGIG6801 |
| 2517205.4 | 397 | 7 | 7 | Candidatus Methanomethylophilaceae archaeon strain CSBr16_184 |
| 2517205.73 | 400 | 7 | 7 | Candidatus Methanomethylophilaceae archaeon B5_69.fa |
| 2517205.17 | 407 | 7 | 7 | Candidatus Methanomethylophilaceae archaeon strain RGIG2195 |
| 2517205.29 | 408 | 7 | 7 | Candidatus Methanomethylophilaceae archaeon strain RGIG5104 |
| 2517205.24 | 409 | 5 | 5 | Candidatus Methanomethylophilaceae archaeon strain RGIG3795 |
| 2774294.4 | 412 | 7 | 7 | Candidatus Methanomethylophilus sp. strain RGIG45 |
| 2517205.11 | 415 | 7 | 7 | Candidatus Methanomethylophilaceae archaeon strain RUG94 |
| 2517205.54 | 418 | 7 | 7 | Candidatus Methanomethylophilaceae archaeon strain RGIG8325 |
| 2517205.46 | 420 | 7 | 7 | Candidatus Methanomethylophilaceae archaeon strain RGIG6905 |
| 2517205.41 | 425 | 6 | 6 | Candidatus Methanomethylophilaceae archaeon strain RGIG6589 |
| 2517205.44 | 427 | 7 | 7 | Candidatus Methanomethylophilaceae archaeon strain RGIG6941 |
| 2517205.22 | 430 | 6 | 6 | Candidatus Methanomethylophilaceae archaeon strain RGIG3602 |
| 2517205.72 | 431 | 7 | 7 | Candidatus Methanomethylophilaceae archaeon strain 10 |
| 2517205.18 | 431 | 7 | 7 | Candidatus Methanomethylophilaceae archaeon strain RGIG1591 |
| 2517203.9 | 432 | 7 | 7 | Candidatus Methanomethylophilaceae GG03_MAG_00002 |
| 2517205.58 | 434 | 4 | 4 | Candidatus Methanomethylophilaceae archaeon strain RGIG8065 |

| | | | | |
|---|---|---|---|---|
| 2517205.15 | 438 | 6 | 6 | Candidatus Methanomethylophilaceae archaeon strain MAG098 |
| 2517205.74 | 440 | 7 | 7 | Candidatus Methanomethylophilaceae archaeon B45_maxbin.030.fa |
| 2517205.35 | 441 | 7 | 7 | Candidatus Methanomethylophilaceae archaeon strain RGIG5541 |
| 2517205.14 | 441 | 7 | 7 | Candidatus Methanomethylophilaceae archaeon strain MAG135 |
| 2517205.34 | 444 | 7 | 7 | Candidatus Methanomethylophilaceae archaeon strain RGIG5548 |
| 2517205.12 | 444 | 7 | 7 | Candidatus Methanomethylophilaceae archaeon strain MAG014 |
| 2774294.5 | 445 | 7 | 7 | Candidatus Methanomethylophilus sp. strain RGIG22 |
| 2517205.38 | 445 | 7 | 7 | Candidatus Methanomethylophilaceae archaeon strain RGIG6046 |
| 2774294.7 | 454 | 7 | 7 | Candidatus Methanomethylophilus sp. strain RGIG1463 |
| 2517205.48 | 455 | 7 | 7 | Candidatus Methanomethylophilaceae archaeon strain RGIG6862 |
| 2517205.51 | 455 | 7 | 7 | Candidatus Methanomethylophilaceae archaeon strain RGIG7034 |
| 2517205.6 | 456 | 7 | 7 | Candidatus Methanomethylophilaceae archaeon strain CSSed162cmB_238 |
| 2517205.43 | 458 | 7 | 7 | Candidatus Methanomethylophilaceae archaeon strain RGIG6752 |
| 2517205.42 | 467 | 7 | 7 | Candidatus Methanomethylophilaceae archaeon strain RGIG6539 |
| 2517205.26 | 467 | 7 | 7 | Candidatus Methanomethylophilaceae archaeon strain RGIG4496 |
| 2517205.32 | 473 | 7 | 7 | Candidatus Methanomethylophilaceae archaeon strain RGIG5314 |
| 2517205.37 | 473 | 7 | 7 | Candidatus Methanomethylophilaceae archaeon strain RGIG5942 |
| 2517205.45 | 475 | 7 | 7 | Candidatus Methanomethylophilaceae archaeon strain RGIG6915 |
| 2517205.25 | 475 | 7 | 7 | Candidatus Methanomethylophilaceae archaeon strain RGIG4274 |
| 2774294.11 | 480 | 7 | 7 | Candidatus Methanomethylophilus sp. strain RGIG8234 |
| 2517203.18 | 484 | 7 | 7 | Candidatus Methanomethylophilaceae GG10_MAG_00009 |
| 2517205.13 | 486 | 7 | 7 | Candidatus Methanomethylophilaceae archaeon strain MAG196 |
| 2517205.47 | 488 | 7 | 7 | Candidatus Methanomethylophilaceae archaeon strain RGIG6908 |
| 2774294.13 | 488 | 7 | 7 | Candidatus Methanomethylophilus sp. strain RGIG8423 |
| 2517205.66 | 488 | 7 | 7 | Candidatus Methanomethylophilaceae archaeon strain RGIG9098 |
| 2517205.39 | 492 | 7 | 7 | Candidatus Methanomethylophilaceae archaeon strain RGIG6768 |
| 2774294.6 | 496 | 7 | 7 | Candidatus Methanomethylophilus sp. strain RGIG1506 |
| 2517205.16 | 500 | 7 | 7 | Candidatus Methanomethylophilaceae archaeon strain RGIG1513 |
| 2517205.36 | 501 | 7 | 7 | Candidatus Methanomethylophilaceae archaeon strain RGIG5987 |
| 2517205.27 | 502 | 7 | 7 | Candidatus Methanomethylophilaceae archaeon strain RGIG4717 |
| 2774294.12 | 536 | 7 | 7 | Candidatus Methanomethylophilus sp. strain RGIG8455 |
| 2517203.24 | 543 | 7 | 7 | Candidatus Methanomethylophilaceae PG05_MAG_00045 |
| 1291539.8 | 1128 | 7 | 7 | Candidatus Methanomethylophilus PG09_MAG_00127 |
| 2774294.15 | 1155 | 7 | 7 | Candidatus Methanomethylophilus sp. UW_MP_METH1_3 |
| 1291540.7 | 1163 | 7 | 7 | Candidatus Methanomethylophilus alvus strain MAG221 |
| 2774294.14 | 1187 | 6 | 6 | Candidatus Methanomethylophilus sp. B13_60.fa |
| 1915608.3 | 1208 | 7 | 7 | Candidatus Methanomethylophilus sp. UBA78 |
| 1908203.3 | 1237 | 7 | 7 | uncultured Candidatus Methanomethylophilus sp. strain RUG779 |
| 1769296.4 | 1294 | 7 | 7 | Candidatus Methanomethylophilus sp. 1R26 |
| 1908203.4 | 1308 | 7 | 7 | uncultured Candidatus Methanomethylophilus sp. strain hRUG898 |
| 1291540.5 | 1308 | 7 | 7 | Candidatus Methanomethylophilus alvus strain 271 |
| 1291540.8 | 1350 | 7 | 7 | Candidatus Methanomethylophilus alvus strain ERR321646-bin.11 |
| 1291540.4 | 1448 | 7 | 7 | Candidatus Methanomethylophilus alvus strain Mx-05 |
| 1291540.6 | 1462 | 7 | 7 | Candidatus Methanomethylophilus alvus strain MGYG-HGUT-02456 |
| 1236689.6 | 1504 | 7 | 7 | Candidatus Methanomethylophilus alvus Mx1201 |
| 2774294.9 | 0 | 0 | 0 | Deleted |

#### Gene Family Statistics

Gene families are ranked by alignment score combining mean per-position variability, alignment length, and gappiness.

| PGFam | Align. Score | Align. Length | Num Seqs | Mean Sqr Freq | Prop Gaps | Used In Analysis | Product |
|---|---|---|---|---|---|---|---|
| PGF_00421879 | 15.08 | 699 | 88 | 0.571 | 0.127 | True | DNA topoisomerase VI subunit B (EC 5.99.1.3) |
| PGF_00020796 | 13.58 | 578 | 93 | 0.565 | 0.146 | True | Methyl coenzyme M reductase system component A2 |

| | | | | | | | |
|---|---|---|---|---|---|---|---|
| PGF_00421761 | 13.52 | 319 | 89 | 0.757 | 0.078 | True | DNA repair and recombination protein RadA |
| PGF_00013509 | 12.83 | 606 | 89 | 0.521 | 0.242 | True | IMP cyclohydrolase (EC 3.5.4.10) / Phosphoribosylaminoimidazolecarboxamide formyltransferase (EC 2.1.2.3) |
| PGF_00007012 | 11.66 | 502 | 90 | 0.521 | 0.269 | True | GTP-binding and nucleic acid-binding protein YchF |
| PGF_00020791 | 10.25 | 210 | 93 | 0.708 | 0.088 | True | Methyl coenzyme M reductase operon protein C |
| PGF_00049827 | 9.85 | 311 | 89 | 0.559 | 0.185 | True | SSU rRNA (adenine(1518)-N(6)/adenine(1519)-N(6))-dimethyltransferase (EC 2.1.1.182) |

### Strategies to Increase Single-Copy Gene Number

Number of single-copy genes (7) was less than requested (100).
Examining the number of genes per genome in the 'Genome Statistics' table above may indicate incomplete or plasmid entries with few genes which may be removed.
Criteria for calling single copy genes can be made more lenient by increasing Max Allowed Deletions and/or Max Allowed Duplications.

Omitting one of the following sets of genomes will provide the approximate boost in single-copy genes:

| scGene Boost | Genomes to Omit |
|---|---|
| 1 | 2517205.21 2517205.22 2517205.28 2517205.30 2517205.40 2517205.49 2517205.5 2517205.53 2517205.56 2517205.59 2517205.61 2517205.64 2517205.7 |
| 1 | 2517203.17 2517205.28 2517205.31 2517205.32 2517205.49 2517205.57 2517205.58 2517205.60 2517205.65 2517205.67 2517205.71 2774294.10 2774294.3 2774294.4 2774294.5 2774294.7 2774294.8 |
| 1 | 2517205.13 2517205.19 2517205.3 2517205.30 2517205.41 2517205.5 2517205.59 2517205.62 2517205.63 2517205.7 2517205.9 2774294.10 |
| 1 | 2517205.41 2517205.53 2517205.60 2517205.64 2517205.65 2517205.68 2517205.69 2517205.7 2517205.71 |
| 1 | 2517205.11 2517205.13 2517205.19 2517205.22 2517205.28 2517205.3 2517205.35 2517205.37 2517205.4 2517205.51 2517205.52 2517205.54 2517205.55 2517205.57 2517205.59 2517205.60 2517205.62 2517205.8 2774294.10 2774294.3 2774294.7 |
